# Functional diversity among sensory neurons from efficient coding principles

**DOI:** 10.1101/131946

**Authors:** Julijana Gjorgjieva, Markus Meister, Haim Sompolinsky

## Abstract

In many sensory systems the neural signal is coded by the coordinated response of heterogeneous populations of neurons. What computational benefit does this diversity confer on information processing? We derive an efficient coding framework assuming that neurons have evolved to communicate signals optimally given natural stimulus statistics and metabolic constraints. Incorporating nonlinearities and realistic noise, we study optimal population coding of the same sensory variable using two measures: maximizing the mutual information between stimuli and responses, and minimizing the error incurred by the optimal linear decoder of responses. Our theory is applied to a commonly observed splitting of sensory neurons into ON and OFF that signal stimulus increases or decreases, and to populations of monotonically increasing responses of the same type, ON. Depending on the optimality measure, we make different predictions about how to optimally split a population into ON and OFF, and how to allocate the firing thresholds of individual neurons given realistic stimulus distributions and noise, which accord with certain biases observed experimentally.

## Introduction

The efficient coding hypothesis states that sensory systems have evolved to optimally transmit information about the natural world given limitations on their biophysical components and constraints on energy use [1]. This theory has been applied successfully to explain the structure of neuronal receptive fields in the mammalian retina [2, 3] and fly lamina [4, 5] based on the statistics of natural scenes. Similar arguments have been made to explain why early sensory pathways often split into parallel channels that represent different stimulus variables, for example different auditory waveforms [6], or local visual patterns [7]. Even neurons that encode the same sensory feature often split further into distinct types. One such commonly encountered diversification is into ON and OFF types: ON cells fire when the stimulus increases and OFF cells when it decreases. This basic ON-OFF dichotomy is found in many modalities, including vertebrate vision [8], invertebrate vision [9], thermosensation [10], and chemosensation [11]. Furthermore, among the neurons that encode the same sensory variable with the same sign, one often encounters distinct types that have different response thresholds, for example, among touch receptors [12] and electroreceptors [13]. The same principle seems to apply several synapses downstream from the receptors [14], and even in the organization of the motor periphery, where motor neurons that activate the same muscle have a broad range of response thresholds [15]. In the present article we consider this sensory response diversification among neurons that represent the same variable and explore whether it can be understood based on a nonlinear version of efficient population coding.

One reason why the ON and OFF pathways have evolved may be to optimize information about both increments and decrements in stimulus intensity by providing excitatory signals for both [16]. For instance, if there were only one ON neuron, then such a cell would need high baseline firing rate to encode stimulus decrements, which can be very costly. We, and others have previously addressed the benefits for having ON and OFF cells in a small population of just two cells [17–19]. However, it remains unclear how a population of many neurons could resolve this issue by tuning their thresholds so that they jointly code for the stimulus. Since ON and OFF neurons often exhibit a broad distribution of firing thresholds [12–15], an important question is thus, what distribution of thresholds yields the most efficient coding. Here we study optimal information transmission in sensory populations comprised of different mixtures of ON and OFF neurons, including purely homogeneous populations with neurons of only one type, e.g. ON, that code for a common stimulus variable by diversifying their thresholds.

Traditionally, efficient population coding has either optimized linear features in the presence of noise [2, 3, 20,21], or nonlinear processing in the limit of no noise or infinitely large populations [22–25]. We simultaneously incorporate neuronal nonlinearities and realistic noise at the spiking output, which have important consequences in finite populations of neurons, as encountered biologically. We develop the problem parametrically in the neuronal noise and the distribution of stimuli that the cells encode, allowing us to make general predictions applicable to different sensory systems.

What quantity might neural populations optimize? We consider two alternative measures of optimal coding that are in common use [22, 26–30]: first we maximize the mutual information between stimulus and response without any assumptions about how this information should be decoded, and second we optimize the estimate of the stimulus obtained by a linear decoder of the response. The two criteria lead to different predictions both on the optimal ON/OFF ratio and the distribution of optimal thresholds. When constraining the maximal firing rate of each cell, we find that counter to our expectations the mutual information is identical for any mixture of ON and OFF cells once the thresholds of all cells are optimized. This result is independent of the shape of the stimulus distribution and the level of neuronal noise. However, the total mean spike count is the lowest for the population with equal numbers of ON and OFF cells, making this arrangement optimal in terms of bits per spike. Optimizing the linear decoder requires determining not only the cells’ thresholds, but also the decoding weights in order to minimize the mean square error between the stimulus and its estimate. Under this criterion, the optimal ON/OFF mixture and cells’ thresholds depend on the asymmetries in the stimulus distribution and the noise level, and can account for certain biases observed experimentally in different sensory systems. We also make distinct predictions for the optimal distribution of thresholds under the two optimality measures, noise level and stimulus distributions, providing insight into the diverse coding strategies of these populations across different sensory modalities and species where these differences are encountered.

## Results

### Population coding model

We develop a theoretical framework to derive the coding efficiency and response properties of a population of sensory neurons representing a common stimulus (Fig. 1A). We specifically consider populations with responses of opposite polarity, ON and OFF, which increase or decrease their response as a function of the common sensory variable; thus, our theory applies to any sensory system where ON and OFF pathways have been observed, for example, heat-activated and cold-activated ion channels in thermosensation [31, 32], mechanosensory neurons [33, 34], or retinal ganglion cells which code for the same spatial location and visual feature with different thresholds [18]. As a special case, we consider populations of neurons with a single polarity, which increase their response as a function of the common sensory variable, for example, olfactory receptor neurons that code for the same odor at different concentrations [35–37]. Each model neuron encodes information about a common scalar stimulus through the spike count observed during a short coding window. The duration of this coding window, *T*, is chosen based on the observed dynamics of neuronal responses, which is typically in the range of 10-50 ms [28, 38]. Neuronal spike counts are stochastic and their mean is modulated by the stimulus through a discrete response function with a finite number of responses. Discretization in neural circuits occurs on many levels [39]; for example, previous experimental studies have found that sensory neurons use discrete firing rate levels to represent continuous stimuli [4, 27, 40]. Furthermore, theoretical work has shown that the optimal neuronal response functions are discrete under different measures of efficiency [27, 41–43].

**Figure 1.**
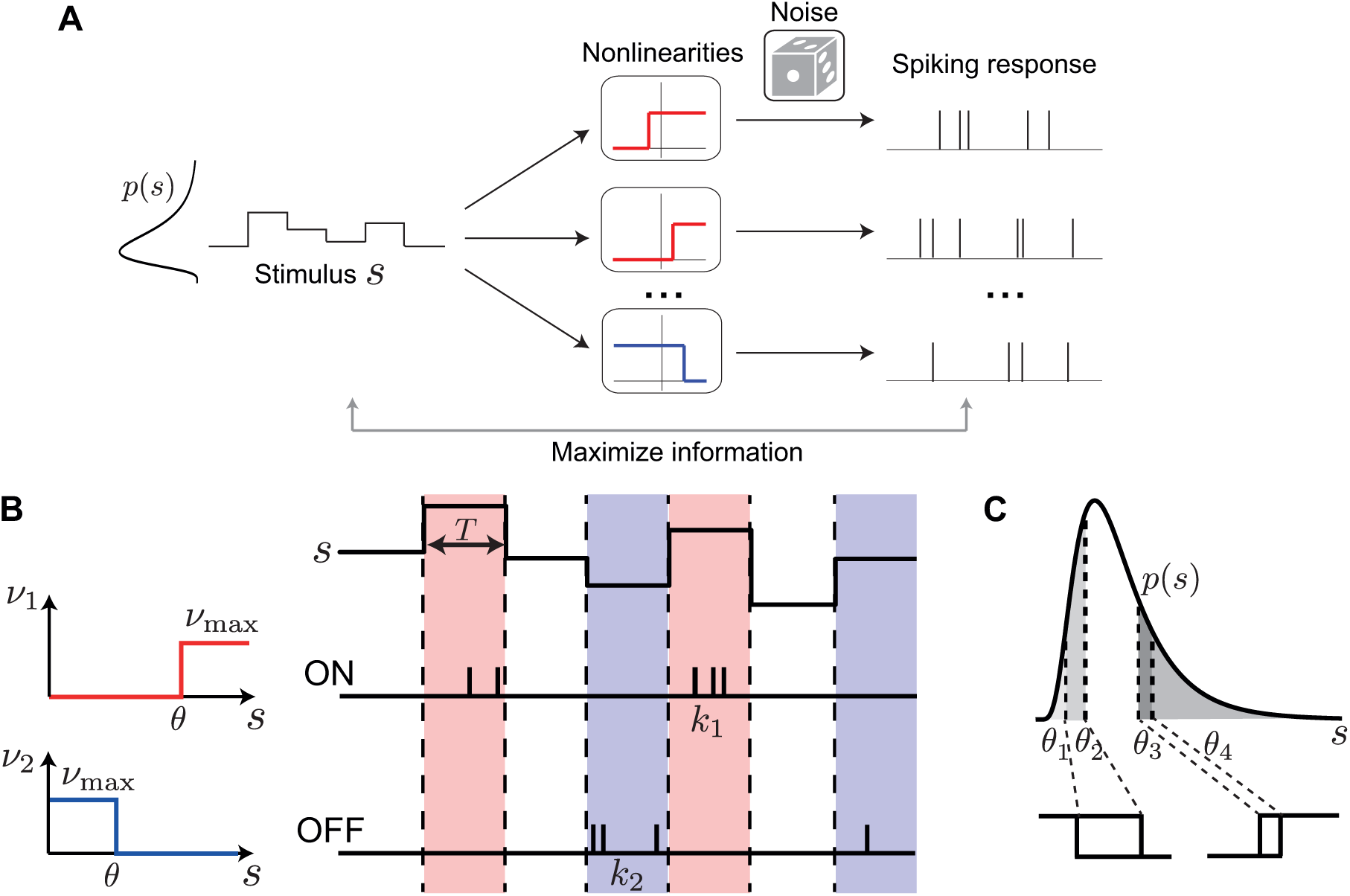
Neuron model and population coding framework. **A**. Framework schematic. A stimulus *s* from a probability distribution *p*(*s*) is encoded by the spiking responses of a population of ON (red) and OFF (blue) cells. We optimize the cells’ nonlinearities by maximizing the mutual information between stimulus and spiking response. **B**. Each cell is described by a binary response nonlinearity *ν* with a threshold *θ* and maximal firing rate *ν*_max_. During a coding window of fixed duration *T* the stimulus is constant and the spike count *k* is drawn from a Poisson distribution with a mean rate *ν*. **C**. When measuring coding efficiency using the mutual information between stimulus and spike count response, the neurons’ thresholds can be interpreted as quantiles of the original stimulus distribution, thus mapping an arbitrary stimulus distribution *p*(*s*) into a uniform distribution (four thresholds shown).

The best way to discretize a neural signal depends on many factors, including noise, stimulus statistics and biophysical constraints [39]. Under the constraint of short coding windows encountered in many sensory areas, optimizing a single response function results in a discretization with two response levels, i.e. a binary response function [27, 28, 41–43]. Binary response functions also offer a reasonable approximation of neural behavior in several systems [28, 44, 45]. Therefore, we assumed that ON (OFF) neurons fire Poisson spikes with an average mean count *ν*_max_ whenever the stimulus intensity is above (below) their threshold *θ*_*i*_, and zero otherwise, i.e. *ν*_*i*_(*s*) = *ν*_max_Θ(*s* − *θ*_*i*_) for ON neurons and *ν*_*i*_(*s*) = *ν*_max_Θ(*θ*_*i*_ − *s*) for OFF neurons (Fig. 1B), where Θ is the Heaviside function.

### Maximal mutual information for mixtures of ON and OFF neurons

What should be the number of ON vs. OFF cells and the distribution of their firing thresholds in a population of neurons that optimally represent a given stimulus? To answer these questions, we first maximize the Shannon mutual information between stimulus and population response, in search of a simple efficient coding principle that could explain ON-OFF splitting and, more generally, threshold diversification. We perform the optimization while constraining the expected spike count *R* = *ν*_max_*T* for each cell. Biophysically, such a constraint on the maximal firing rate arises naturally from refractoriness of the spike-generating membrane. We have analytically proven the following theorem (see Methods):

#### Equal Coding Theorem

For a population of any number *N* of ON and OFF Poisson neurons coding a one-dimensional stimulus in a fixed time window *T* by binary rate functions with maximal firing rate *ν*_max_, the mutual information is identical for all ON/OFF mixtures when the thresholds are optimized, for all *N*, *ν*_max_ and stimulus distributions.

Specifically, the maximal information is achieved in the case when the ON and OFF cells do not overlap, so that all ON thresholds are bigger than all OFF thresholds. For example, consider a mixed population of ON and OFF cells. To calculate the information conveyed by this entire population, we imagine first observing only the ON cells, and in a second step the remaining OFF cells. If one of the ON cells fired a spike, we know the stimulus is in that cell’s response range, and therefore we do not learn additional information from observing the OFF cells. If none of the ON cells fired, we gain additional information from observing the OFF cells. One can make the same argument if the remaining cells are all ON cells, or indeed any other mixture. Careful consideration shows that the maximal information gained from that remaining cell population is the same whether they are ON cells or OFF cells (see Methods and S1 Text). Hence, the homogeneous and any mixed ON-OFF population achieve the same maximal information.

The value of this maximal information depends on the expected spike count, *R* = *ν*_max_*T*. We introduce the parameter *q* = *e*^−*R*^, which ranges from *q* = 0 in the noiseless limit of high firing rate, and *q* = 1 in the high noise limit of low firing rate. We show that for any ON-OFF mixture (including the homogeneous with only ON cells), the maximal information achieved with optimized thresholds is (Fig. 2A, see Methods)

**Figure 2.**
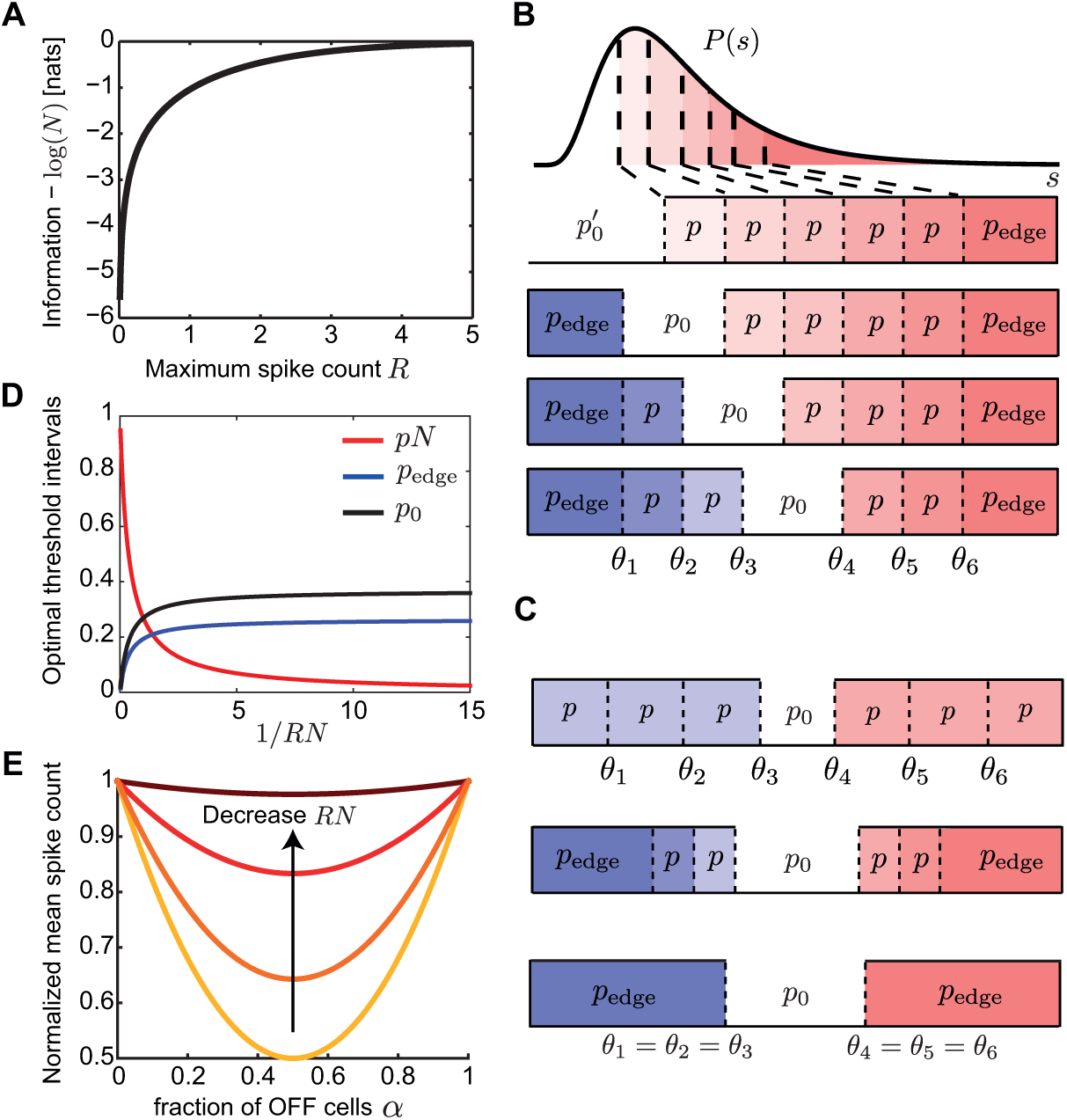
Mutual information when constraining the expected spike count. **A**. The mutual information between stimulus and response for any mixture of *N* ON and OFF cells is identical when constraining the expected spike count, *R*. **B**. The optimal threshold intervals for all possible mixtures of ON (red) and OFF (blue) cells in a population of *N* = 6 cells that achieve the same mutual information about a stimulus from an arbitrary distribution *p*(*s*). **C**. The optimal threshold intervals for the equal ON-OFF mixture in a population of *N* = 6 cells and different values of *R* (equivalently, noise); see also D. Top: low noise (*RN* → *∞*); middle: intermediate noise (*RN* = 1); bottom: high noise (*RN* → 0). **D**. The optimal threshold intervals as a function of 1*/RN*. **E**. The mean spike count required to transmit the same information (see A) by populations with a different fraction of OFF cells (*α*), normalized by the mean spike count of the homogeneous population with *α* = 0. The different curves denote *RN* = {0.1, 1, 5, 100}.

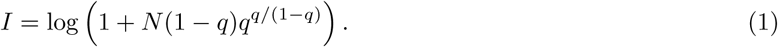

We further ensured that the conclusion of equal coding holds in a population of two cells independent of the Poisson noise model we assumed, which has zero noise when the the firing rate of a neuron is zero. Specifically, we investigated information transmission introducing a spontaneous firing rate under the same Poisson model, as well as empirically measured sub-Poisson noise from salamander retinal ganglion cells [17, 28] (see S1 Text).

The above equation 1 allows us to exactly compute the maximal information that would be reached by a population of neurons as a function of the number of neurons *N* and the level of noise *q* assuming optimality, without resorting to expensive numerical calculations [46]. Even if real biological systems do not perform optimally, this quantity could be used as an upper bound for the largest possible information that the system could transmit under the appropriate constraints. In the noiseless limit, *R* →∞ (i.e. *q* = 0), where the neurons are deterministic, *I* reaches its upper bound *I* = log(*N* + 1). The effect of noise is most prominent when *R* is of order 1*/N*, so that the total spike count *RN* is of order 1, implying that the signal-to-noise of the entire population is of order 1. We call this the **high noise regime**, and here we obtain *I* → log(*RN/e* + 1), where *e* denotes exp(1).

### Optimal distribution of thresholds

We next asked what distribution of thresholds in the population of ON and OFF cells achieves this maximal mutual information. In the case of a discrete rate function, we can replace *θ*_*i*_ by the corresponding cumulative threshold (fraction of stimuli below threshold), which essentially maps the stimulus distribution into a uniform distribution from 0 to 1 (Fig. 1C). Since the stimulus dependence enters only through these values, the maximal mutual information is independent of the stimulus distribution *p*(*s*), provided that the stimulus cumulative distribution is continuous. Instead, the information depends on the areas of *p*(*s*) between consecutive thresholds. It is therefore useful to define the optimal threshold intervals 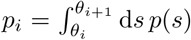 where the neurons’ threshold are ordered *θ*_1_ ≤ … *≤ θ*_*N*_ (and we define the special *θ*_0_ = −*∞* and *θ*_*N*+1_ = *∞*). We find a surprisingly simple structure for the optimal *p*_*i*_ (Fig. 2B,C). The optimal thresholds divide stimulus space into intervals of equal area, which depend on the noise level, *q*,

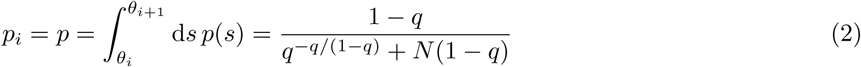

for all *i, except* for the two *‘edge’* intervals,

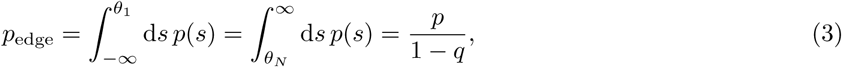

and the *‘silent’* interval that separates the ON and OFF thresholds, *p*_0_ = 1 − (*N* − 2)*p* − 2*p*_edge_ (see Methods). Note that for the homogeneous population,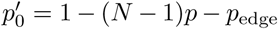. We call this optimal threshold structure the **infomax** solution.

We consider several limiting cases: first, a large population *N* ≫ 1 and maximal firing rate per neuron *R*, which is much larger than 1*/N*, i.e. 1 − *q* = *𝒪*(1). We call this the **large population regime**. In this regime, *p*_edge_ = *p* = *p*_0_ = 1*/*(*N* + 1), so the *N* thresholds divide stimulus space into *N* + 1 equal intervals (Fig. 2C top, D). In this large population regime, we can rewrite the optimal thresholds as a continuous function of cumulative stimulus space; we replace *θ*_*i*_ with *θ*(*x*), where *x* = *i/N* is the threshold index between 0 and 1. Then the optimal thresholds equalize the area under the stimulus density,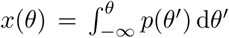. Therefore, the population of cells achieves ‘histogram equalization’ in that it uses all the available response symbols at equal frequency, as has been shown before for a single cell with many discrete signaling levels in the limit of no noise [4, 47].

In contrast, when the noise is high so that *RN* → 0, the system performs redundant coding so that each *p*_*i*_ is infinitesimally small and the only two substantial threshold intervals are the edge intervals *p*_edge_ = 1*/e*, and the silent interval, *p*_0_ = 1 − 2*/e*, which separates the ON and OFF thresholds (Fig. 2C bottom, D). This *p*_0_ is the only non-noisy response state where the firing rate of each cell is zero. This implies that the optimal solution is to place all ON thresholds at roughly the same value, and similarly all OFF thresholds at another value (Fig. 2C bottom). This solution maximizes redundancy across neurons in the interest of noise reduction [2, 48, 49], consistent with various experimental and theoretical work [2, 48, 49]. Interestingly, for a small population of two cells, we (and others) have previously shown that in the presence of additional input noise before the signal passes through the nonlinearity, this ‘redundant coding’ regime exists for larger range of noise values [17–19].

In summary, we have derived the total mutual information and distribution of optimal thresholds in a population of binary neurons coding for the same stimulus variable for any stimulus distribution and noise level. While our results agree with previous work in the limit of no noise and an infinite population, we make unique and novel predictions – notably the surprisingly regular structure of the threshold intervals and invariance in information transmission for any ON/OFF mixture – in populations of any number of neurons and with sizable noise relevant for majority sensory systems.

### Optimizing information per spike predicts equal ON/OFF mixtures

Our analysis so far showed that maximizing the information equally favors all ON-OFF mixtures independent of the noise level, although the exact distribution of population thresholds at which this information is achieved depends on noise. However, different sensory systems show dominance of OFF [50], dominance of ON [34, 51], or similar numbers of ON and OFF [34]. Therefore, we next explored what other criteria might be relevant for neural systems under the efficient coding framework. We considered that neural systems might not just be optimized to encode as much stimulus information as possible, but might do so while minimizing metabolic cost. Therefore, for each ON-OFF mixed population achieving the same total information (Fig. 2A), we calculated the mean spike count used to achieve this information. In the large population regime, if *α* denotes the fraction of OFF cells, the mean spike count per neuron is *r*(*α*) = *R*(*α*^2^ + (1 − *α*)^2^)*/*2 (Methods). This mean spike count per neuron is minimized at *α* = 1*/*2, where it is half of the mean spike count for the homogeneous population, *r*(0) = *R/*2 (Fig. 2E). This implies that it is most efficient to split the population into an equal number of ON and OFF cells. As the noise increases, the relative benefits of the equally mixed relative to the homogeneous population decrease (Fig. 2E). In the high noise regime, all mixtures produce roughly the same mean spike count per neuron of *R/e* (Fig. 2E). Therefore, if a sensory system is optimized to transmit maximal information at the lowest spike cost, our theory predicts similar numbers of ON and OFF neurons, which is consistent with ON-OFF mixtures encountered in some sensory systems [34].

### Minimizing mean square error of the optimal linear readout

The efficient coding framework does not specify which quantity neural systems optimize to derive their structure. Until now, we have used the mutual information as a measure of coding efficiency because it tells us how well the population represents the stimulus without regard for how it can be decoded. An alternative criterion for coding efficiency is the ability of downstream neurons to decode this information. A simple biologically plausible decoding mechanism, commonly used in previous studies, is linear decoding [22, 26, 29, 30, 52]. Does this alternative measure of efficiency generate the same predictions for how sensory populations coding for the same stimulus variable should allocate their resources to ON vs. OFF neurons? Here, we examine the accuracy of a downstream neuron that estimates the stimulus value *s* using a weighted sum of spike counts *n*_*i*_ of the upstream population of neurons with thresholds *θ*_*i*_ (Fig. 3A)

**Figure 3.**
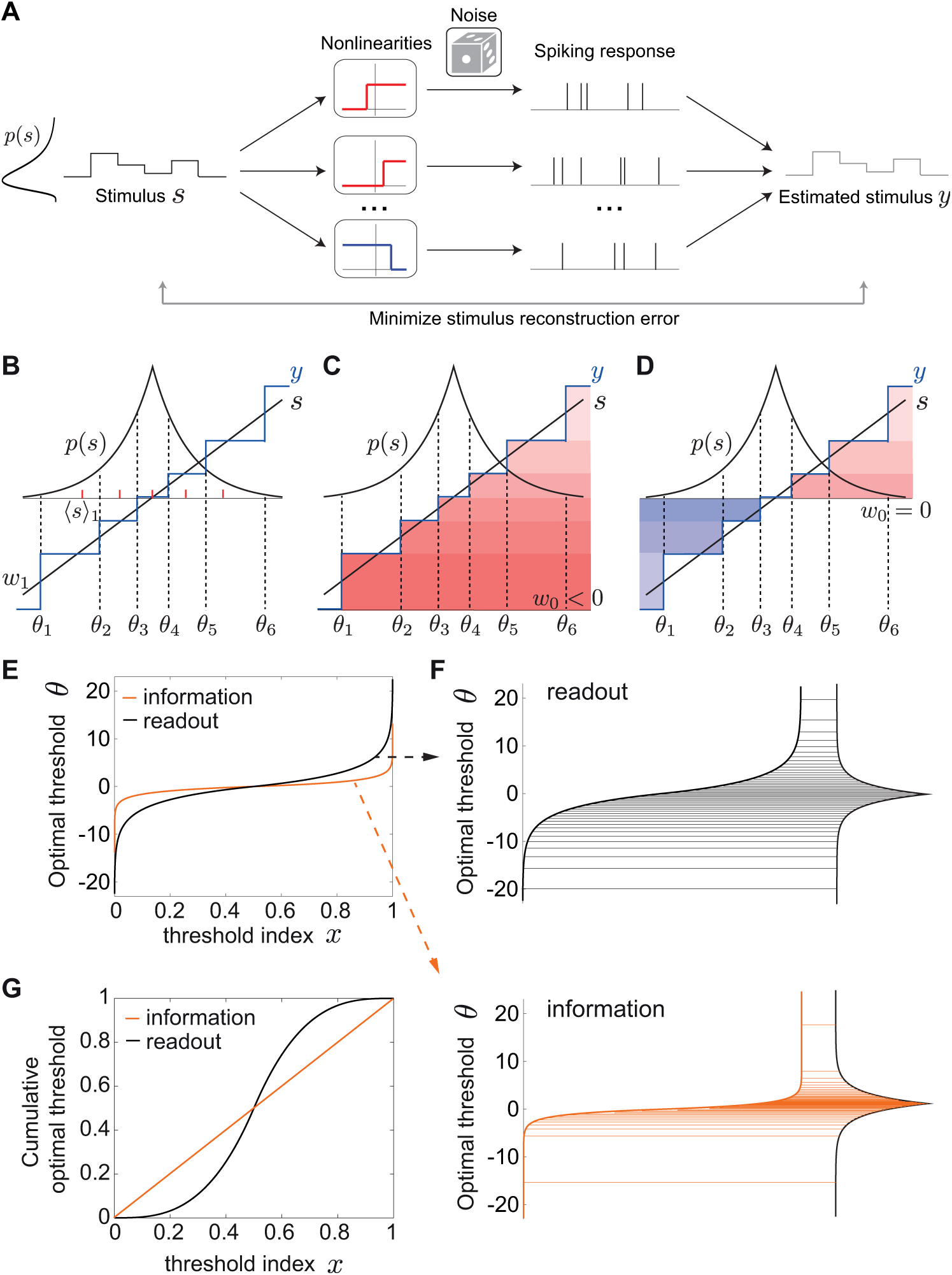
Optimal linear decoding of stimuli. **A**. Framework schematic. A stimulus *s* from a probability distribution *p*(*s*) is encoded by the spiking responses of a population of ON (red) and OFF (blue) cells. We optimize the cells’ nonlinearities by minimizing the mean squared error (MSE) between the original stimulus *s* and the linearly reconstructed stimulus *y* from the spiking response. **B**. Minimizing the MSE between a stimulus *s* (black) and its linear estimate *y* (blue) by a population of (6) ON and OFF cells, in the absence of noise. We show the optimal weight *w*1 and the center of mass ⟨*s*⟩1 of the first threshold interval (red dashes). **C**,**D**. Any ON-OFF population can achieve the same error with the same set of optimal thresholds and weights but a different constant, *w*0. **C**. 6 ON cells (*w*0 < 0). **D**. 3 OFF and 3 ON cells (*w*0 = 0). **E**. The optimal thresholds equalize not the area under the stimulus density (as in the case of the mutual information), but the area under its one-third power (Eq. 8). The optimal thresholds are shown for the Laplace distribution. **F**. The information maximizing thresholds partition the Laplace distribution into intervals that code for stimuli with higher likelihood of occurrence (bottom), while minimizing the MSE pushes thresholds to favor rarer stimuli near the tails of the distribution (top). Threshold distributions are the same as in E. **G**. The cumulative optimal thresholds 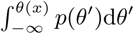 (compare to E).

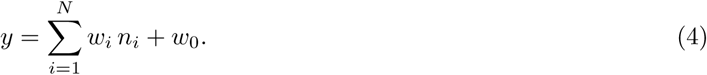

The weights *w*_*i*_, constant *w*_0_ and thresholds *θ*_*i*_ are optimized to minimize the mean square estimation error (MSE).

### Accuracy of the optimal linear readout without noise

We first consider the scenario of low noise (*q* → 0, or equivalently, *R* → *∞*), in which case the limitation on the accuracy of the stimulus reconstruction comes solely from the discreteness of the rate functions of each cell in the population (Fig. 3B). Unlike maximizing the information, when minimizing the MSE both weights and thresholds depend on the stimulus distribution *p*(*s*) (Fig. 3).

Interestingly, we find that in this low noise limit, the optimal MSE is proportional to 1*/N* ^2^ and is the same for all ON/OFF mixtures, including the homogeneous population with all cells of the same type (Fig. 3C,D; see Methods). The optimal decoding weights are given by

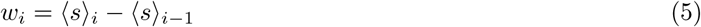

where ⟨*s*⟩_*i*_ are the centers of mass of intervals of *p*(*s*) intersected by neighboring thresholds

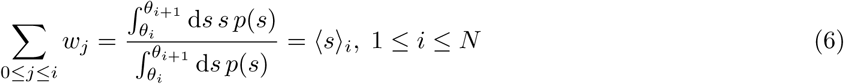

where we have defined *θ*_*N*+1_ = *∞*. The optimal thresholds are the average of two neighboring centers of mass

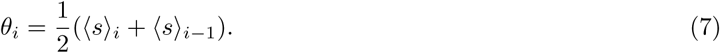

The constant term and the stimulus interval not coded by any cell depend on the ON/OFF mixture (Fig. 3C,D; Methods). This gives a recursive relationship that from a set of initial thresholds converges to the optimal solution (see Methods).

To see how this threshold distribution is different than the one predicted by the mutual information, we first consider the large population regime. As for the mutual information, we can rewrite the thresholds *θ*_*i*_ as a continuous function *θ*(*x*) of the cumulative stimulus values *x* = *i/N* between 0 and 1. Interestingly, the optimal thresholds equalize not the area under the stimulus density, as in the case of the mutual information, but the area under its one-third power

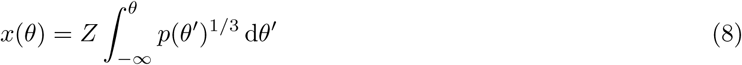

where *Z* is a normalization factor. This result has been previously derived in the context of minimizing the distortion introduced in a pulse-coupled-modulation system due to quantization [53] (reviewed in [54]), as well as in the context of neural coding which maximizes the *L*_*p*_ reconstruction error of the maximum likelihood decoder, of which the mean squared error is the special case for *p* = 2 [25].

We invert the relationship in equation 8 to derive the optimal thresholds *θ*(*x*). Since the optimal MSE depends on the stimulus distribution, from now on we consider the Laplace distribution *p*(*s*) = 1*/*2 *e*^−|*s*|^, which arises when evaluating natural stimulus distributions [23, 55] and has a higher level of sparseness than the Gaussian distribution. In this case, the optimal thresholds become (Fig. 3E,F; see Methods):

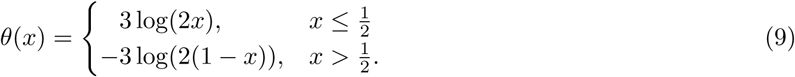

The thresholds derived from maximizing information are the same except that the pre-factor is 1 instead of 3, making them less spread out in the tails (Fig. 3F). In particular, the largest thresholds (in magnitude) are ±3 log(2*N*) when optimizing the MSE, three times as large as in the infomax case, ±log(2*N*). To highlight the different predictions for the optimal thresholds under the two efficiency measures, we also plot the cumulative optimal thresholds 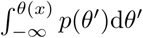 (Fig. 3G). While the optimal strategy when maximizing the information is to emphasize stimuli with higher likelihood of occurring, minimizing the MSE of the optimal linear readout pushes thresholds logarithmically towards relatively rare stimuli near the tails of the stimulus distribution (Fig. 3F,G).

Taken together, we conclude that in the absence of noise our theory derives equal performance of all ON and OFF mixtures under the two optimality criteria, information maximization and minimizing the optimal linear readout. However, a key difference between the two criteria is the theoretically predicted optimal distribution of thresholds.

### Mixed ON/OFF populations in the presence of noise

In biologically realistic scenarios with non-negligible noise, however, we find that mixed ON/OFF populations show a dramatic improvement of the MSE over predominantly homogeneous populations (Fig. 4A). For the Laplace distribution we have considered so far, and different noise values, we find that the optimal fraction of OFF cells in the population is *α* = 1*/*2. Although there is a unique best ON/OFF mixture, the best linear stimulus reconstruction achieved by other populations with ON-OFF mixtures closer to the optimal 1/2 mixture is similar (i.e. the MSE around *α* = 1*/*2 is flat). The worst stimulus reconstruction is achieved by the homogeneous population with all cells of ones type (all ON or all OFF), which has the highest MSE. As the noise *q* decreases (*R* increases) further, this difference in performance between the mixed and homogeneous populations becomes quite dramatic, see for example *R* = 1 (Fig. 4A).

**Figure 4.**
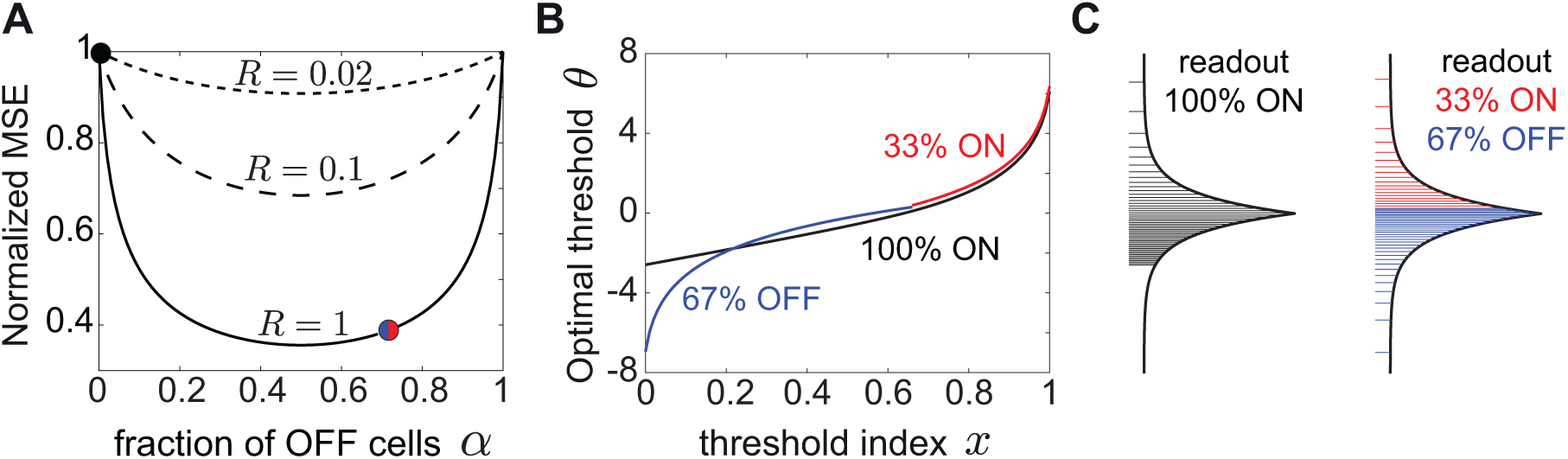
Optimal linear decoding of stimuli with noise depends on the ON/OFF mixture. **A**. The MSE as a function of the fraction of OFF cells in the population, *α*, for a different expected spike count, *R*. The MSE was normalized to the MSE for the homogeneous population of all ON cells. The MSE is shown for *N* = 100 cells and for the Laplace distribution. Symbols indicate the MSE values realized with the thresholds in B and C. **B**. The optimal thresholds for the homogeneous population (black) partition the Laplace stimulus distribution starting with a much larger first threshold than the mixed population with 2/3 OFF cells (blue) and 1/3 ON cells (red). **C**. The optimal thresholds for the Laplace distribution for a homogeneous population (black) and a mixed population with 2/3 OFF cells (blue) and 1/3 ON cells (red). In B and C, *R* = 1. Note the difference in the optimal threshold distribution between the mixed ON/OFF and the homogeneous ON population, especially for small *x* = *i/N* (logarithmic in blue vs. linear in black).

In addition to the big difference in coding performance between mixed and homogeneous populations, incorporating biologically realistic noise also affects the theoretically derived distribution of optimal thresholds (Fig. 4B). While in mixed populations the thresholds are distributed logarithmically towards relatively rare values at the tails of the stimulus distribution (Eq. 9; see Fig. 4B,C and Methods), for the homogeneous population the optimal thresholds exhibit a distinct asymmetry. A large fraction of thresholds are distributed linearly as a function of their index, while the remaining thresholds are distributed logarithmically as before:

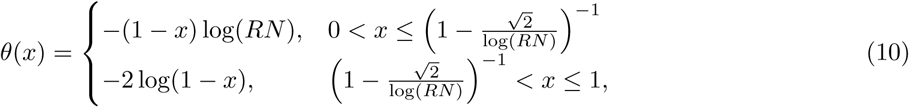

although the noise has the effect of concentrating the thresholds near more likely stimuli, increasing the redundancy of the code. Moreover, the smallest threshold for the homogeneous population is much larger than the smallest threshold for any mixed population, suggesting that there is a large region of stimuli that is not coded by any cell in the homogeneous case (Fig. 4C), which is the reason for the significantly lower MSE.

In summary, using the MSE of the optimal linear decoder as a measure of efficiency can fundamentally alter our conclusions about how to split a population into ON and OFF cells and how to distribute the population thresholds to achieve the optimal stimulus reconstruction. At biologically realistic noise levels, coding by mixed ON-OFF populations is much better than by a homogeneous population, with qualitatively distinct optimal threshold distributions.

### The optimal ON-OFF mixture of the linear readout depends on the asymmetry in the stimulus distribution

Since the MSE as a measure of efficiency depends on the stimulus distribution, we asked how the stimulus distribution can affect optimal population coding. The distribution of natural stimuli may be asymmetric around the most likely stimulus. For example, the distribution of contrasts in natural images, and the intensity of natural sounds are indeed skewed towards more negative values [20, 56–61]. Therefore, we instead consider an asymmetric Laplace distribution *p*(*s*) ∝ *e*^*s/τ*^− for *s* < 0 and *p*(*s*) ∝ *e*^−*s/τ*^+ for *s* ≥ 0 where we take *τ*_−_ *> τ*_+_. Minimizing the MSE one finds that the optimal way to divide a population into ON and OFF respects these stimulus asymmetries. Increasing the negative stimulus bias *τ*_−_*/τ*_+_ favors more OFF cells (Fig. 5A,B). The optimal thresholds for these different stimulus biases are best compared in the cumulative space of stimulus (Fig. 5C). Increasing the bias also pushes the thresholds towards more negative stimulus values, which occur with higher probability than positive stimuli.

**Figure 5.**
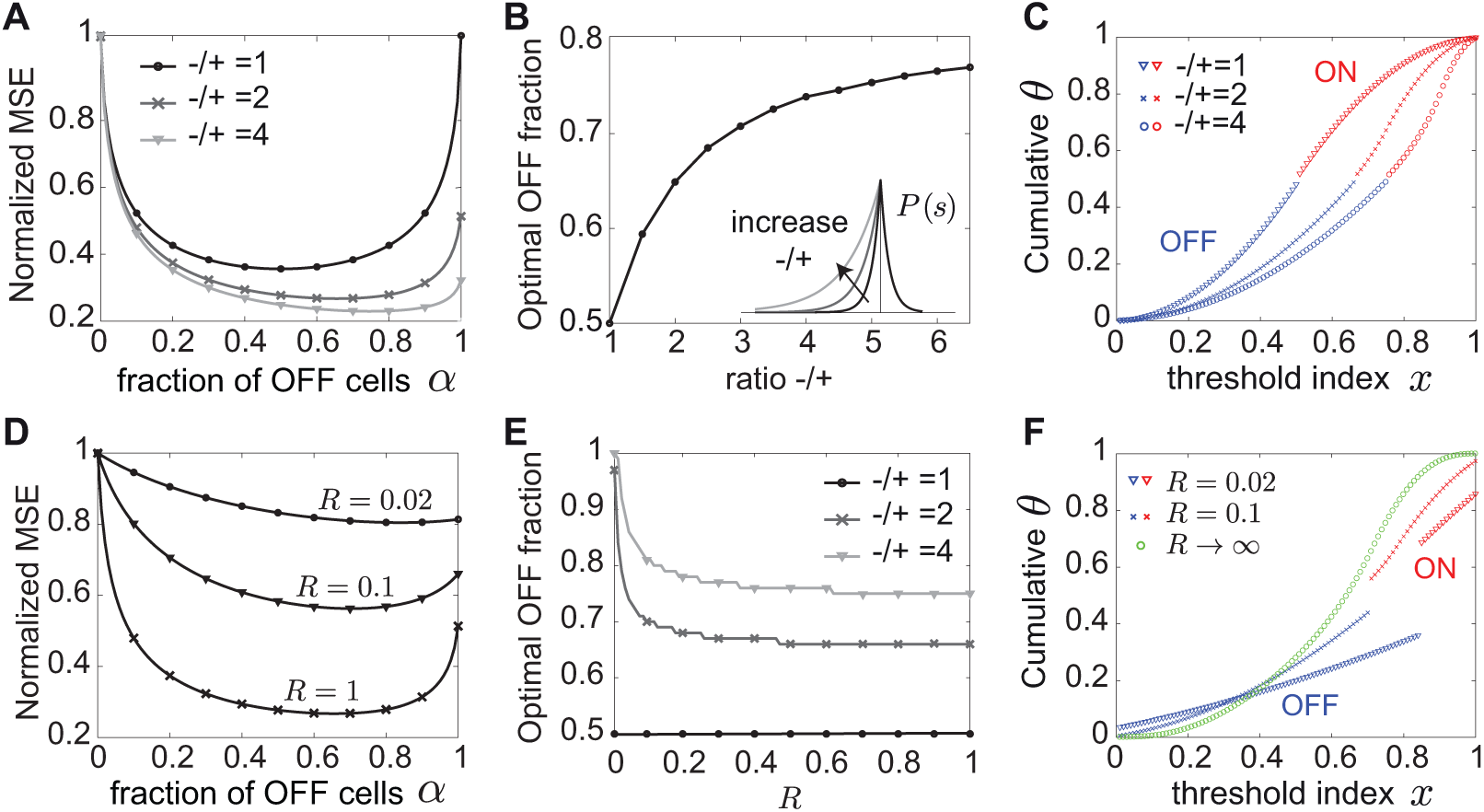
The optimal ON/OFF mixture derived from the linear readout is tuned to asymmetries in the stimulus distribution. **A**. The MSE as a function of the fraction of OFF cells (*α*) normalized to that for the homogeneous population of all ON cells (*α* = 0). The MSE is shown for an asymmetric Laplace distribution with varying negative to positive bias −*/*+, expected spike count *R* = 1 and *N* = 100 neurons. **B**. The optimal fraction of OFF cells as a function of stimulus bias of the asymmetric Laplace distribution and *R* = 1. **C**. The optimal thresholds for the ON-OFF mixtures (50%, 66% and 75%) in A that yield the lowest MSE, while varying negative to positive bias −*/*+ = {1, 2, 4}. **D**. Same as A but for an asymmetric Laplace distribution with a negative bias −*/*+ = 2 and varying *R* (equivalently, noise). **E**. The optimal fraction of OFF cells as a function of *R* for different stimulus bias of the asymmetric Laplace distribution. **F**. The optimal thresholds for the ON-OFF mixtures (84%, 70% and *any*) in D that yield the lowest MSE, while varying *R* = {0.02, 0.1, *∞*}.

At a fixed level of stimulus bias, increasing the noise further accentuates the asymmetry in the optimal ON-OFF mixture (Fig. 5D,E). As the noise becomes non-negligible, the optimal thresholds lose the logarithmic spread at the tails of the stimulus distribution and begin to code for more likely stimuli that occur with a higher probability. At the same time, a larger region of stimulus values near the median is no longer coded by any cells, i.e. the gap between ON and OFF thresholds becomes larger (Fig. 5F). Had we considered the limit of zero noise or infinitely large populations as previous studies [22–25], we would not have been able to identify these differences between the optimal thresholds that result in conditions of biologically realistic noise and finite populations.

In summary, our theory predicts different optimal ON-OFF numbers at which the lowest MSE is achieved depending on asymmetries in the stimulus distribution and the noise level. Indeed in nature, the relative predominance of ON and OFF cells in diverse sensory systems can be different (Table 1). Therefore, if we know the natural stimulus distribution being encoded by a population and the bounds on cells’ firing rates, we can predict the optimal ON and OFF numbers, as well as the response thresholds of the cells and compare them to experimental observations.

**Table 1.**
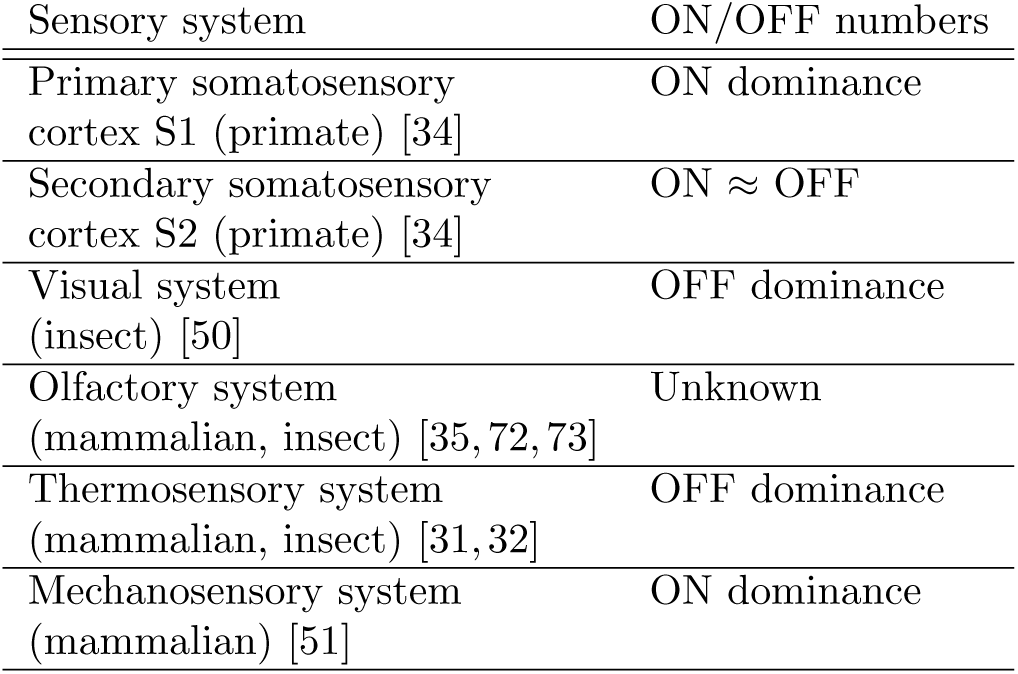
List of experimentally measured ON and OFF neuron numbers in different sensory systems.

| Sensory system | ON/OFF numbers |
| --- | --- |
| Primary somatosensory cortex S1 (primate) [34] | ON dominance |
| Secondary somatosensory cortex S2 (primate) [34] | ON ≈ OFF |
| Visual system (insect) [50] | OFF dominance |
| Olfactory system (mammalian, insect) [35, 72, 73] | Unknown |
| Thermosensory system (mammalian, insect) [31, 32] | OFF dominance |
| Mechanosensory system (mammalian) [51] | ON dominance |

### Predicting stimulus distributions from experimentally measured thresholds

Here we propose to reverse our efficient coding framework and starting from an experimentally measured distribution of thresholds, to predict the distribution of natural stimuli that the thresholds could be optimized to encode (Fig. 6A). This could be particularly relevant for sensory systems where the distribution of the sensory variable being encoded is unknown. We decided to test this approach on odor concentration coding in the olfactory system of *Drosophila* larvae given recently published data [37]. The first stage of olfactory processing in *Drosophila* larvae is implemented by a population 21 olfactory receptor neurons (ORNs), which code for a broad space of odorants and concentrations [37]. We hypothesized that these ORNs might have distributed their thresholds at different concentrations to optimally encode any particular odor. In the classical efficient coding approach, knowing the distribution of odor concentrations would allow us to predict the optimal thresholds. In the reversed approach that we use here, knowing the distribution of thresholds allows us to predict the distribution of concentrations of a known odor (Fig. 6A).

**Figure 6.**
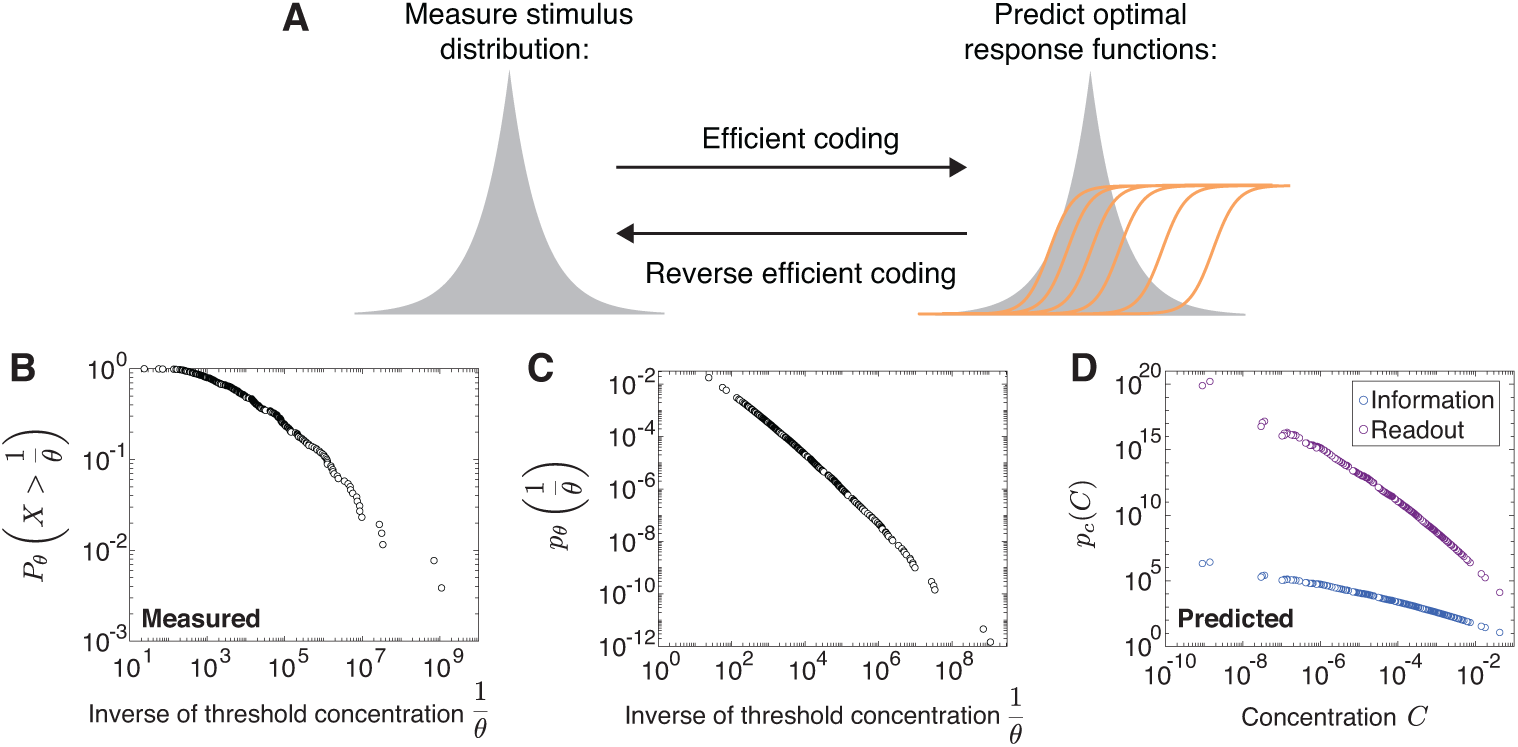
Deriving a distribution of stimulus intensities from experimentally measured thresholds. **A**. Our efficient coding framework enables us to predict the optimal distribution of thresholds given a known stimulus distribution. By reversing our framework, we derive the stimulus distribution from a distribution of measured thresholds assuming optimal coding under the two optimality criteria. **B**. Log-log plot of the cumulative distribution of the inverse of thresholds from measured dose-response curves of the entire population of ORNs in the *Drosophila* larva olfactory system [37]. This is well described by a power law with exponent −0.42. **C**. The probability distribution of the inverse of optimal thresholds derived from the data in B. This is well described by a power law with exponent −0.58. **D**. Predicted distribution of concentrations across different odorants when assuming optimal coding by maximizing information or minimizing the MSE of the best linear decoder. This is well described by a power law with exponents −0.58 and −1.74, respectively. The proportionality constant is not shown.

A recent study estimated these thresholds by recording from the entire ORN population [37]. The responses for 34 odorants over a five-fold magnitude in concentration were well described by a common Hill function with a shared steepness, but different activation thresholds. Pooling all thresholds across the different odorants and concentrations revealed a power law distribution. To use this threshold distribution in our theoretical framework, where a range of thresholds codes for the intensity of a single stimulus, we had to make a critical assumption. Specifically, we assumed that the population thresholds spanning the range of concentrations for any one odor are a shuffled version of the population thresholds for other odorants. This was justified by an analysis of a related data set [35], in which the distribution of ORN firing rates was found to be stereotyped across different odors [36].

Using our optimal coding framework with a population of only ON neurons (since ORNs have monotonically increasing response functions with concentration), we derived the mostly likely stimulus distribution of odorant concentrations for each of the two efficiency measures. The predicted distribution of odorant concentrations follows a power law distribution with an exponent determined by the efficiency measure. Given a measured distribution of thresholds which follows a power law with an exponent of −0.58 (Methods, Fig. 6B,C) and assuming an infomax code we predict that the distribution of odorant concentrations should also be a power law with an exponent of −0.58 (Methods, Fig. 6D). In contrast, assuming a code that minimizes the stimulus reconstruction error, the distribution of odorant concentrations should be a power law with an exponent of − 1.74 (Methods, Fig. 6D). Indeed, many processes like convection and turbulence can generate power law dynamics [62], but the exact exponents will need to be determined, for instance by measuring the volatiles from natural environments [63, 64]. Although complex temporal dynamics in the stimulus can further complicate ORN coding of fluctuating odorant concentrations, the measured temporal filter across ORNs is remarkably stereotyped, suggesting that the olfactory code is similar between static and dynamic odor environments.

We note that in our analysis we explicitly assume that the goal of the olfactory system is to estimate the concentration of any one odor with high fidelity, therefore it is only valid for experiments where only one odor is present. However, the optimization problem faced by the olfactory system might be different, i.e. to determine which, of many, odors are present. Therefore, it is possible that the optimal thresholds in these two cases may be different.

## Discussion

Information in neural circuits is processed by many different cell types, but it remains a challenge to understand how these distinct cell types work together. Here we treat a puzzling aspect of neural coding, how do discrete cell types conspire to collectively encode a single relevant variable in responses of opposite polarity? To evaluate such a population code we built on the framework of efficient coding and extended it in several novel ways: by considering nonlinear processing, biologically realistic levels of noise, short coding windows, and the coordination of responses in populations of any size – factors which may vary across sensory systems. We then derived two aspects of the population code, namely how to optimally split a population into ON and OFF cells, and how to allocate the thresholds of the individual neurons as a function of the noise level, the stimulus distribution and the optimality measure.

### Optimal ON-OFF mixtures and comparison to experimental data

We considered two different measures of coding efficiency that are in common use [22, 26–30]: the mutual information between stimulus and responses, and the mean squared error of the linearly reconstructed stimulus. The first aspect of our predictions applies to the expected mixture of ON and OFF cells. If one chooses mutual information as the efficiency measure, then all ON/OFF mixtures in the population perform identically once the thresholds are adjusted (Fig. 2). This result holds independent of the noise level and the shape of the stimulus distribution, and generalizes for response functions with any number of discrete firing rate levels. However, the number of spikes required for this performance, and thus the metabolic cost, differs greatly depending on the ON/OFF ratio. If one considers the information per spike as the relevant measure, then a system with equal number of ON and OFF cells is most efficient.

When we require the stimulus to be read out by an optimal linear readout, different ON/OFF mixtures also achieve similar coding performance but only in the absence of noise (Fig. 3). In the biologically relevant regimes of non-negligible noise, noise has a dramatic influence on the optimal performance realized by different ON/OFF mixtures (Fig. 4). Populations with a similar number of ON and OFF cells have a much smaller decoding error than populations dominated by one cell type. The extreme case of the homogeneous population performs substantially worse that any mixed population (Fig. 4). In the case of asymmetries in the stimulus distribution, as encountered in many natural sensory stimulus distributions [20, 56–61], minimizing the linear reconstruction error predicts that the optimal ON/OFF mixture should be tuned to these asymmetries and the amount of noise (Fig. 5).

How do these predictions accord with known neural codes? Since our theory applies to populations of sensory neurons that code for the same stimulus variable, we need to consider sensory systems where this is the case. Analyzing raw stimulus values, such as the light intensity in a natural scene or the intensity of natural sounds, results in distributions which are skewed towards negative stimuli [20, 56–61]. Our linear decoding theory then predicts that more resources should be spent on OFF. Indeed, in the fly visual system, the OFF pathway is overrepresented in the circuit for computations that extract motion vision, with the L1 neurons being responsible for the processing of ON signals, while both L2 and L3 neurons for OFF [9, 50]. These neurons are repeated in each cartridge, thus together code for the same spatial location. Hence, at least for the fly visual system, our efficient coding results are in accord with naturally encountered ON/OFF ratios. In contrast, the vertebrate retina represents a visual stimulus with spikes across diverse types of retinal ganglion cells, which differ in their spatial and temporal processing characteristics [65, 66]. Certain types of ganglion cell come in ‘paramorphic pairs,’ meaning an ON-type and an OFF-type that are similar in all other aspects of their visual coding. A previous study by Ratliff et al. (2010) derived the optimal numbers of ON and OFF retinal ganglion cells for encoding natural scenes assuming maximal information transmission, as a function of the spatial statistics in these natural stimuli. In their model, every ganglion cell in the population encodes a different stimulus variable, because it looks at a different spatial location. In contrast, our theory can only be applied to populations that code for the same same stimulus feature, which may in fact contain only one of each type (ON and OFF), but requires further experiments to determine the exact numbers. To properly account for all thirty types of retinal ganglion cells will require more complete models that include the spatial dimension and the encoding of different visual features.

Besides the visual system, there are other examples in biology where different numbers of ON and OFF cells are encountered, and where our theory more naturally applies with populations of neurons encoding a one-dimensional stimulus (Table 1). Single neurons in monkey somatosensory cortex show diverse ON and OFF responses to the temporal input frequency of mechanical vibration of their fingertips. While most neurons in primary somatosensory cortex (S1) tune with a positive slope to the input frequency (ON), about half of the neurons in secondary somatosensory cortex (S2) behave in the opposite way (OFF) [33, 34]. Opposite polarity pathways are also observed in thermosensation, where receptor proteins activated directly by positive and negative changes in temperature enable the detection of thermal stimuli. Four mammalian heat-activated and two cold-activated ion channels have been shown to function as temperature receptors [31, 32]. Given these observations of ON-OFF asymmetries, one is led to conclude that information per spike may not be the cost function that drove evolution of this system, since that would predict equal numbers of ON and OFF cells. Thus, whether these different experimental observations are consistent with maximizing mutual information, optimal linear decoding, or yet a different objective function or task (e.g. [67–69]), remains to be seen (see also our discussion on the generality of assumptions). Other neuronal systems are candidates for similar analysis, for instance, auditory nerve fibers [44], motor cortex [70], and primary vestibular neurons [71].

### Optimal threshold distributions and comparison to experimental data

Beyond predicting ON-OFF numbers, which has been the main focus of different models about the vertebrate retina [20], we also predict the structure of response thresholds. Generally, maximizing information implements an optimal strategy which emphasizes stimuli that occur with higher probability (Fig. 3, 4). In the limit of low noise, this is consistent with the well-known strategy of ‘histogram equalization’ [4], but we generalize this result to any amount of biologically realistic noise. Importantly, the optimal interval size depends on the level of noise with larger noise favoring smaller threshold intervals, implying a strategy closer to redundant coding. In contrast to the information, minimizing the mean square error of the linear readout implements a more conservative strategy that utilizes more cells in the encoding of rarer stimuli due to a larger error penalty(Fig. 3, 4).

Our theoretical framework applies to the case when the distribution of stimuli encoded by the cells is known, and the only problem is to estimate the value of the stimulus by appropriately distributing the cells’ thresholds. In the case of vision, for example, this implies estimating the light intensity or contrast level. A direct test of our theoretical predictions for the optimal thresholds would require simultaneous measurement of the population response thresholds, which is within reach of modern technology [66]. In the meantime, we reversed our theoretical approach and starting from an experimentally measured distribution of thresholds, we predicted the distribution of natural stimuli that the thresholds might optimally encode. We applied this approach for the population of ORNs in the olfactory system of *Drosophila* larvae. However, unlike vision, applying our framework to olfaction presents a different problem. Here, the goal of the olfactory system is primarily to determine whether or not an odor is present, not its concentration. Therefore, our analysis is only appropriate when only one odor is present, and it can be inferred with high certainty. In this case, we assumed that the ORNs code for the distribution of concentrations of the present odor by diversifying their thresholds. The ORNs’ experimentally described tuning curves were identical in shape, with response thresholds following a power law distribution [37]. Since the probability distribution of ORN firing rates is stereotyped across different odors [36], we assumed that the thresholds coding the range of concentrations for any one odor are a shuffled version of the thresholds for other odorants. We derived the stimulus distribution of concentrations for any one tested odor, under the two optimality measures, the maximal mutual information and the minimal error of the best reconstructed stimulus (Fig. 6). This threshold distribution was also a power law with an exponent dependent on the efficiency measure. Whether these distributions correspond to distributions of odorant concentrations found in natural olfactory environments remains to be tested, and techniques for collecting the volatiles from natural encountered odors now exist [63, 64]. These distributions would be strongly influenced by processes like convection and turbulence, which can give rise to power law dynamics [62]. Although these are dynamical variables that fluctuate in time, we propose that the distributions can be build by pooling different aspects of the dynamics over extended time periods. In that context, our theoretical framework would apply to populations which have been adapted to these distributions over those long periods of time. It is possible that when considering a different optimization problem implemented by the olfactory system which aims to determine which, of many, odors are present, a very different distribution of thresholds than those our theory predicts would be optimal.

### Generality of assumptions and relationship to previous work

The two efficiency measures that we have used are entirely agnostic about the content of signal transmission. However, faithful encoding of signals is not the only fitness requirement on a sensory system, for example, some stimuli may have greater semantic value than others. Or, the aim may be to extract task-relevant sensory information as in the case of the Information Bottleneck framework [67, 68], or to achieve optimal inference of behaviorally-relevant properties in dynamic stimulus environments [69]. Other recent approaches, such as Bayesian efficient coding, optimize an arbitrary error function [74]. Since our framework aims to encode a stimulus as best as possible, we propose that it may be most appropriate for early sensory processing, where stimulus representation might be the goal.

The efficient coding hypothesis was originally proposed by Attneave [75] and Barlow [1], who studied deterministic coding, in the absence of noise. Since then, many studies have investigated efficient coding strategies under different conditions. Atick and Redlich introduced noise and demonstrated that efficient coding can be used to explain the center-surround structure of receptive fields of retinal ganglion cells, which changes to center-only structure as the signal-to-noise increases [2, 76]. Including nonlinear processing in the limit of low noise produced Gabor-like filters encountered in the primary visual cortex [13]. However, we now know that already the very first stages of processing in many sensory systems are nonlinear, consist of many parallel pathways and exhibit substantial amount of noise [77] – important aspects of coding that we simultaneously incorporate in our analysis. Our work differs from a previous report on ON and OFF cells in the vertebrate retina which proposed a simplified noise model implemented by assuming a finite number of signaling levels (i.e. firing rates), which does not incorporate spiking [20].

We considered a Poisson noise model of spiking which is commonly used in many studies. Our results are especially relevant in the high noise regime, which corresponds to short coding windows commonly encountered in biology, for instance, a few spikes per coding window [28, 38, 78]. In the low noise regime when the coding window is sufficiently long, or there is a large number of neurons, our results agree with previous studies on infomax and the optimal linear readout [4, 25, 79]. Efficient coding in the high noise regime has previously been examined, but only in terms of the transfer function of a single neuron, which was shown to be binary [27, 43]. We go beyond this work and provide analytical solutions for how a population of neurons should coordinate their response ranges to optimally represent a given stimulus in the realistic regimes of short encoding times.

We used a binary rate function to describe single neuron responses because it gives our problem analytical tractability and it still represents a significant departure from previous efficient coding frameworks based on linear processing [2, 3, 20, 21], long coding windows or infinitely large populations [22–25]. Indeed, discretization in neural circuits is a common phenomenon that is not only relevant for sensory coding, but also for neuropeptide signaling, ion channel distributions and information transmission in genetic networks [39, 80]. Considering more general nonlinearities is currently only tractable with numerical simulations or in the case of optimizing a local efficiency measure, the Fisher information, which may not accurately quantify coding performance in finite size populations or biologically realistic noise (e.g. low firing rates or short coding windows) [25, 49, 81–83].

### Summary

Given the ubiquity of ON/OFF pathway splitting in different sensory modalities and species, our framework provides predictions for the optimal ON/OFF mixtures and the functional diversity of sensory response properties that achieve this optimality in many sensory systems based on the distribution of relevant sensory stimuli, the noise level and the measure of optimality. Our theoretical approach is sufficiently general and is not finetuned to the specifics of any one experimental system. The different predictions that we make depending on the model assumptions could help determine the specific optimality criteria operating in different sensory systems where different ON-OFF mixtures and tuning properties have been observed. Directed experiments to compare the predicted and measured threshold distributions will test whether the efficient coding criteria proposed here are a likely constraint shaping the organization and adaptation of sensory systems.

## Materials and Methods

### Mutual information and proof of the Equal Coding Theorem

First we prove the Equal Coding Theorem for a general population with *N* binary neurons. Without loss of generality we assume that the neurons’ thresholds are:

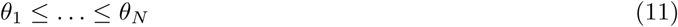

and we define the special *θ*_0_ = −*∞* and *θ*_*N*+1_ = *∞*. The Shannon mutual information between the stimulus *s* and the spiking response ***n*** of the population is the difference between response and noise entropy:

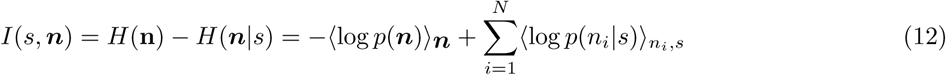

where ⟨*·*⟩_*x*_ denote averages over the distribution *p*(*x*) and *p*(***n***) = ⟨*p*(***n***|*s*) ⟩_*s*_. We assume that stimulus encoding by all neurons is statistically independent conditional on *s* so that

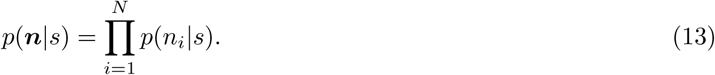

Given the Poisson noise model, knowing the stimulus *s* unambiguously determines the response firing rate *ν*; for instance, for an ON cell if *s* < *θ, ν* = 0 and if *s ≥ θ, ν* = *ν*_max_. We can replace *p*(*n*_*i*_|*s*) with *p*(*n*_*i*_|*ν*) which is Poisson distributed: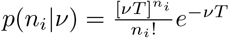.

We prove that *I*(*s*; ***n***) = *I*(***ν, n***). To see this, we write

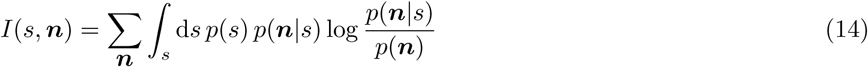

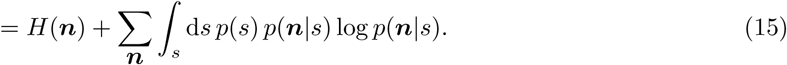

Using Eq. 13, this becomes

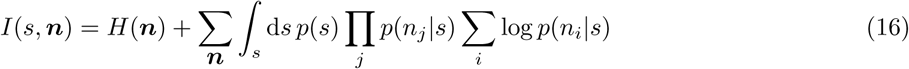

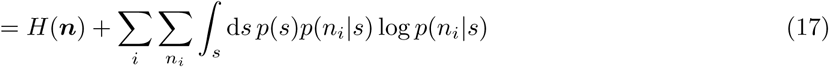

Similarly, we derive

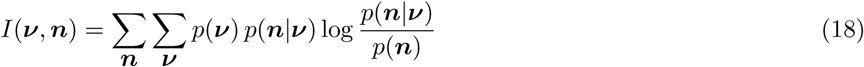

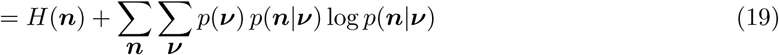

which since

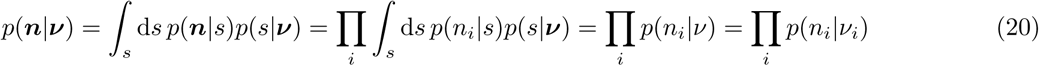

becomes

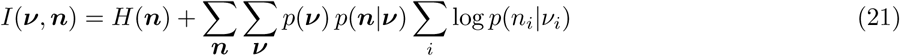

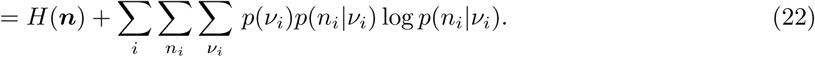

Now, for a given *i* and a corresponding given spike count *n*_*i*_, which without loss of generality we assume is an ON cell with threshold *θ*_*i*_, we take the second term from Eq. 17 and split the integral:

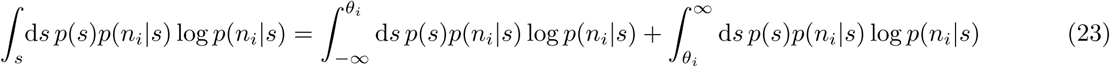

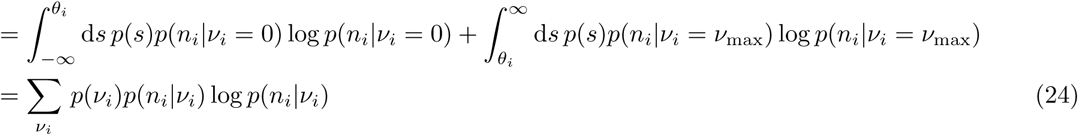

because we can just integrate out the *s*. Therefore, from Eqs. 17, 22 and 24, we get *I*(*s*; ***n***) = *I*(***ν, n***). Note that for a single cell, Nikitin et al. [43] also proved the same equality of information using a different approach.

For a binary response function with two firing rate levels, 0 and *ν*_max_, we can lump together all states with *nonzero* spike counts into a single state which we denote as **1**. Correspondingly, the state with zero spikes is 0. Hence, we can evaluate the mutual information between stimulus and spiking response using the following expressions for the spike count probabilities:

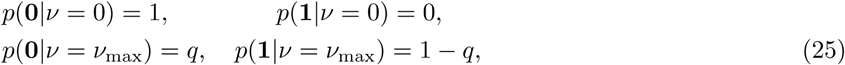

where *q* = *e*^−*R*^ and *R* = *ν*_max_*T* denote the level of noise in the system.

We can derive the expression for the mutual information between stimulus and response given the *N* intervals

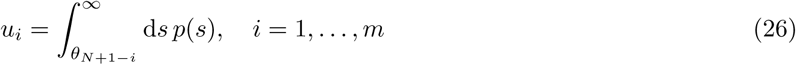

for the *m* ON cells and

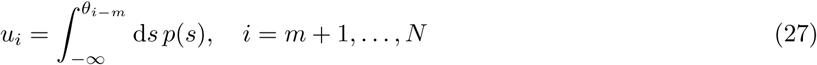

for the OFF cells, see Figure 7.

**Figure 7.**
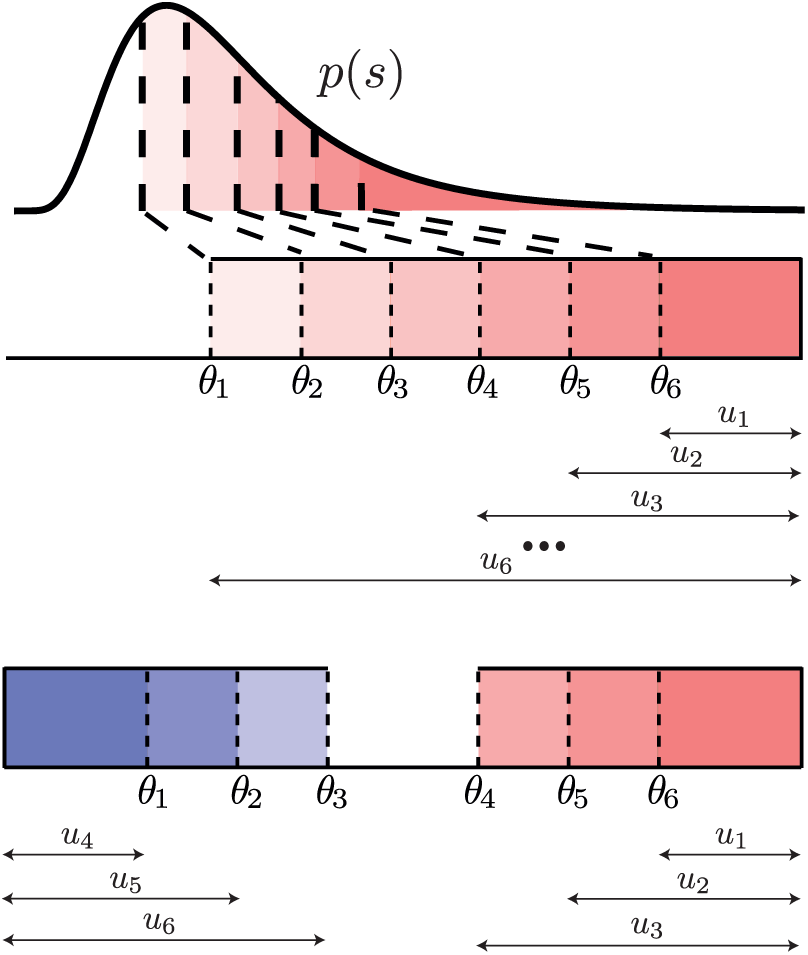
Thresholds *θ*_*i*_ and intervals between thresholds *u*_*i*_ for a population of 6 cells. Top: a homogeneous population with 6 ON cells; bottom: a mixed population with 3 ON and 3 OFF cells.

We prove the Equal Coding Theorem by showing that the mutual information coded by a population of *N* ON cells is the same as that for any arbitrary mixture of ON and OFF cells, for instance, a population with *m* ON cells (with indices 1, 2, … *m*) and *N* − *m* OFF cells (with indices *m* + 1, …, *N*). At the optimal solution the ON cells have larger thresholds than the OFF cells. This is due to the assumed Poisson noise model, where the states at which a given cell’s firing rate is 0 are non-noisy and determine the stimulus with certainty. The total information can be described as the information from observing the ON cells 1, … *m*, plus any additional information gained from observing cells *m* + 1, …, *N*. Below we demonstrate that this additional information is identical independent of whether the *N* − *m* cells are ON (in which case the population is homogeneous and comprised of all ON cells) or OFF type (in which case the population is mixed). This turns out to be the case, as long as the thresholds of the additional *N* − *m* cells are appropriately adjusted.

If a spike was observed from ON cells 1, 2, … *m*, then no additional information is gained from cells *m* + 1, …, *N*, independent of their type because their firing rate is constant over the entire stimulus interval in which cells 1, …, *m* fire. Then, the total mutual information achieved by all *N* cells, *I*_*N*_ (*s*, ***n***), is equal to the mutual information obtained from observing the *m* ON cells, *I*_*m*_(*s*, ***n***):

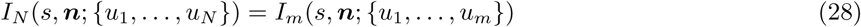

where we explicitly denote the dependence of the mutual information on the threshold intervals, *u*_*i*_’s. If no spike was observed from the ON cells 1, 2, … *m*, then we get additional information from the remaining cells *m* + 1, …, *N*, but we need to consider the change in the stimulus distribution posterior to seeing no spike. Such a change in the stimulus distribution is equivalent to adjusting the thresholds of cells *m* + 1, …, *N*, and as a result, the threshold intervals. If none of the ON cells 1, … *m* fired, then, we can formally write the total information as follows (see Fig. 7):

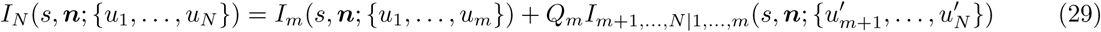

where *Q*_*m*_ is the probability that none of the ON cells 1, … *m* fired, and *I*_*m*+1,…,*N*|1,…,*m*_ is the additional mutual information gained from the remaining *N* − *m* cells with adjusted thresholds, and consequently threshold intervals, 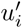.

By writing the information in this manner, we have only assumed that the first *m* cell are ON, but have not assumed anything about the type of the additional *N* − *m* cells. In fact, for any ON-OFF mixture given by the number of ON cells *m*, one can choose the same thresholds *θ*_1_, …, *θ*_*m*_ (and thus thresholds intervals *u*_1_, …, *u*_*m*_) for the first *m* ON cells, and then change the thresholds *θ*_*m*+1_, …, *θ*_*N*_ (and thus threshold intervals *u*_*m*+1_, …, *u*_*N*_) of the remaining *N* − *m* cells so as to produce the same adjusted threshold intervals, 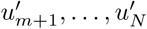.

#### How can this readjustment be done for the different mixtures?

If no spike was observed from the ON cells 1, …, *m*, then the stimulus distribution to be coded by the remaining cells changes from the prior *p*(*s*) to a new posterior distribution

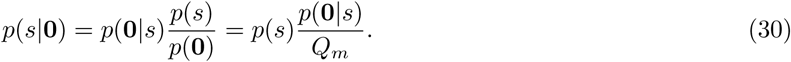

1. **If the remaining** *N* − *m* **cells are ON**, the region of reduced *p*(*s*) is entirely within the response region. Thus, the revised probability of having the stimulus in the response region is

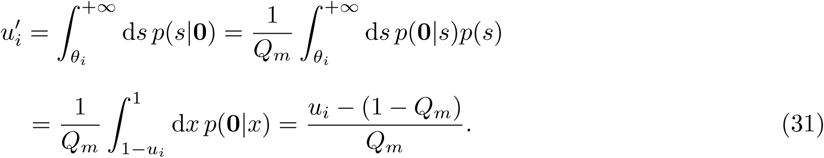

where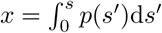.

2. **If the remaining** *N* −*m* **cells are OFF**, the region of reduced *p*(*s*) is entirely outside their response region. Thus, their revised probability is

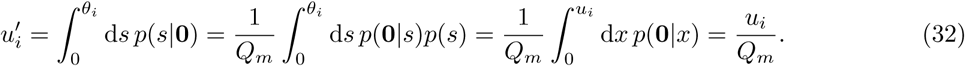

Therefore, the readjustment of the threshold intervals can be done differently for a homogeneous population when the remaining *N* − *m* cells are all ON, vs. a mixed population when the remaining *N* − *m* cels are all OFF. Since *m* can be anything between 1 and *N*, this covers all possible mixtures of ON and OFF cells, where

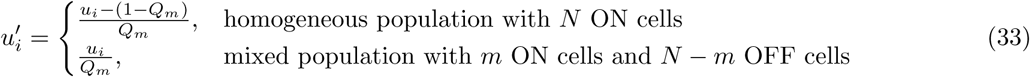

To find the maximal mutual information one needs to maximize Eq. 29 with respect to all the thresholds (i.e. threshold intervals). Since the homogeneous population of *N* ON cells and the mixed population of *m* ON cells and *N* − *m* OFF cells share the same *m* ON cells, maximizing the total mutual information *I*_*N*_ in Eq. 29 is equivalent to maximizing the additional mutual information *I*_*m*+1,…,*N*|1,…,*m*_ gained from the remaining *N* − *m* cells with adjusted threshold intervals according to Eq. 33. This explains why the maximum information is identical between the purely homogeneous population with *N* ON cells and a mixed population where *N* − *m* cells are OFF.

#### Thresholds when optimizing the mutual information: a homogeneous population

Next we derive the optimal thresholds for the homogeneous population with *N* ON cells, and later derive the thresholds of the *N* − *m* OFF cells after swapping.

If the thresholds are ordered in ascending order as assumed above, then *u*_1_ < *u*_2_ < … < *u*_*N*_ (S3 Figure). The mutual information of *N* ON cells can be written as follows. First, for a population of *N* = 1 cells this has the form

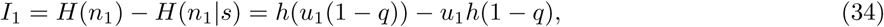

where *h* is the entropy of a binary variable, *h*(*u*) = − *u* log *u* − (1 − *u*) log(1 − *u*). For a population of *N* = 2 cells it has the form

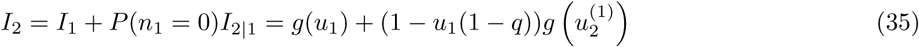

where we have defined *g*(*u*) = *h*(*u*(1 −*q*)) −*uh*(1 −*q*). Here, 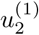 denotes the revised value of *u*_2_ following the observation of cell 1. In general, we use 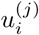 to denote the revised value of *u*_*i*_ after the observation that cell *j* < *i* did not spike. Therefore, for a population of *N* = 3 cells it has the form

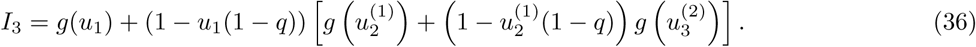

Generalizing this for *N* cells, the information is

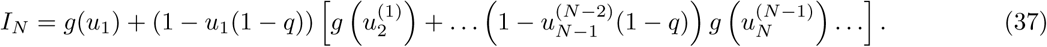

The revised values of 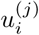 for *i* = 2, … *N* and *j* = 1, … *i* − 1 follow based on readjusting the thresholds depending on the observation of cells 1, … *N* − 1 one at a time. For example, following the observation that cell 1 did not spike, the effective values of *u*_2_, *u*_3_, … *u*_*N*_ are revised to

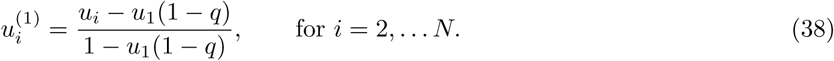

Following the observation that cell 2 did not spike, 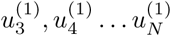 are further revised to

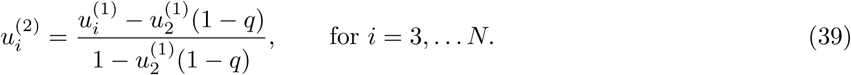

This process continues, until the observation of cell *N* − 1 with the final set of 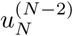 being revised to

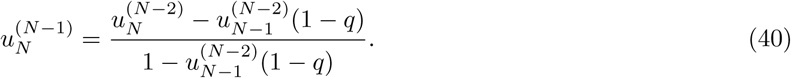

We maximize the information in Eq. 37 with respect to each 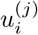. We can do this sequentially: first maximize *I* with respect to 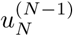, which results in maximizing 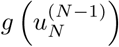. The maximum is obtained at

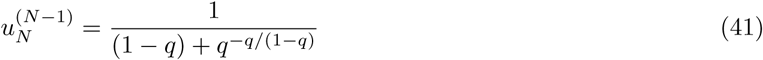

yielding a maximal value of

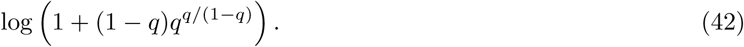

Next, we maximize *I* with respect to 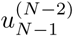, which results in maximizing 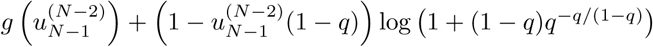. The maximum is obtained at

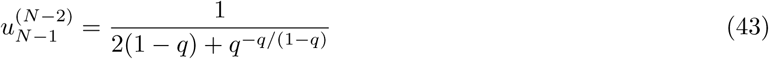

yielding a maximal value of

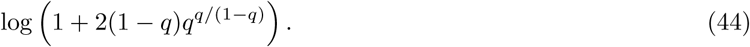

Finally, we maximize *I* with respect to *u*_1_, which results in maximizing *g*(*u*_1_) + (1 −*u*_1_(1 −*q*)) (log 1 + (*N* − 1)(1 −*q*)*q*^−*q/*(1−*q*)^). The maximum is obtained at

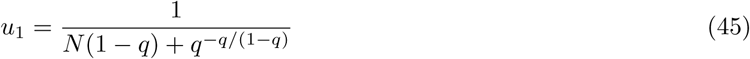

yielding a maximal value of the mutual information as in Eq. 1 in the Results section

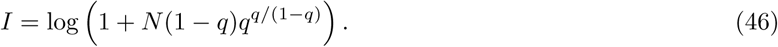

Based on these derivations we can obtain the sequence of

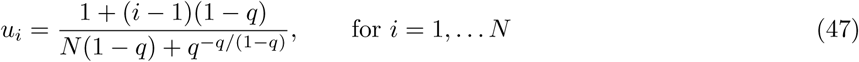

where the difference between consecutive thresholds is given by Eq. 2

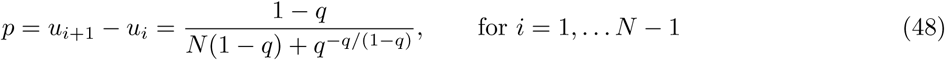

and the ‘edge’ threshold is Eq. 3

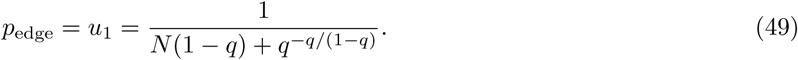

### Thresholds when optimizing the mutual information: a mixed population

With the Equal Coding Theorem we showed that the information for any ON/OFF mixture is the same (Eq. 46). Next, we show how to derive the thresholds for a mixed population since we know that it will have the same mutual information as the homogeneous population. We do this by swapping *N* − *m* of the ON cells into OFF cells, knowing that the thresholds of the ON cells in the new mixed population remain the same, and derive the thresholds for the swapped OFF cells. This means that we need to derive a new set of 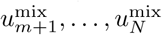 for the OFF population, while keeping *u*_1_, … *u*_*m*_ the same for the ON population. To do this, recall that the thresholds for the OFF cells follow different update rules every time an ON cell is observed (see Eq. 33). In particular,

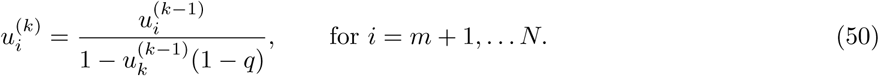

Additionally, following the observation of OFF cell *k* (where *k* = *m* + 1, …, *N* − 1)

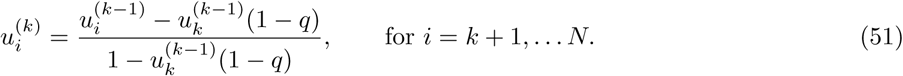

Using these recursions and the values 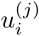 for the ON cells derived previously (Eq. 41 – 45) one can recover the thresholds:

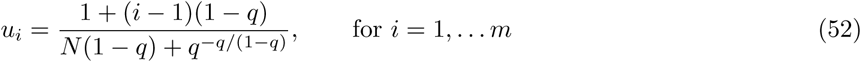

for the ON cells and

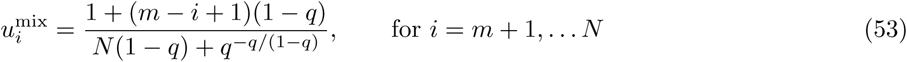

for the OFF cells (in the mixed population case) where the difference between consecutive thresholds (except between the smallest ON and the largest OFF) is given by Eq. 48 and the ‘edge’ thresholds by Eq. 49. From here we can derive the *‘silent’* interval between the smallest ON and the largest OFF that separates the ON and OFF thresholds, *p*_0_ = 1 − (*N* − 2)*p* − 2*p*_edge_.

### Mean firing rate when optimizing the mutual information

Given the optimal thresholds, the mean firing rate per neuron in a population with *m* ON cells is:

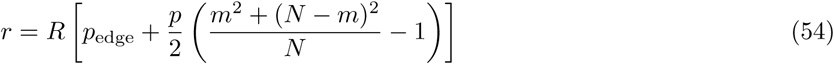

In the large population regime, with *α* = *m/N* the fraction of ON cells, the mean firing rate per neuron is

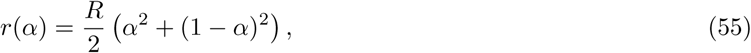

however, in the high noise regime this becomes independent of *α*

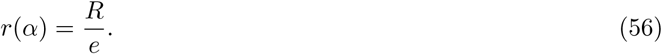

### Optimal linear readout without noise

We present here the derivation for the homogeneous population with only ON cells when *R* → *∞*. The linear stimulus estimate of *s* (Eq. 4) can be written as:

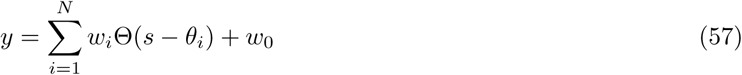

where *w*_*i*_ represent the decoding weights and the responses are given by the binary Heaviside functions with thresholds *θ*_*i*_. Then the mean square error between the original and the estimated stimulus can be written as:

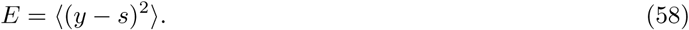

In the case of the homogeneous population, we can emulate the constant term *w*_0_ as the weight of an additional neuron with threshold *θ*_0_ = −*∞*. Then

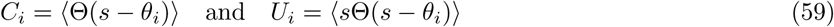

so the error can be written as:

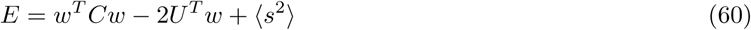

where since ⟨Θ(*s* − *θ*_*i*_)Θ(*s* − *θ*_*j*_)⟩ = ⟨Θ(*s*− max(*θ*_*i*_, *θ*_*j*_)), for *i* ≥ *j*, we can write: *C*_*ij*_ = *C*_*i*_. Optimizing with respect to the weight will gives us

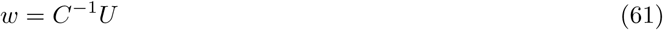

which we can rewrite as (Eq. 6 in the Results section):

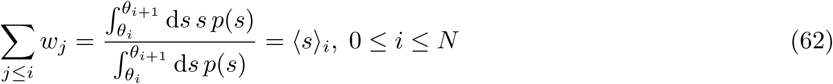

with *θ*_*N*+1_ = ∞ and (Eq. 5 in the Results section):

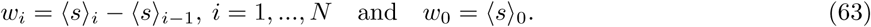

Optimizing with respect to the thresholds:

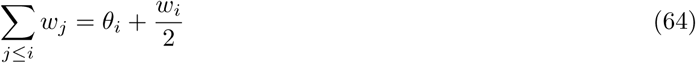

which gives

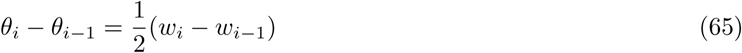

and from this we can derive (Eq. 7 in the Results section):

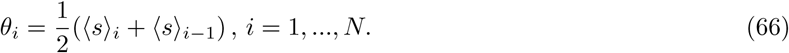

Optimizing with respect to the constant term yields:

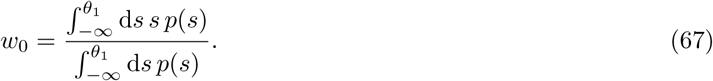

To solve these equations numerically, we implement an iterative procedure that rapidly converges to the optimal solution: starting from an ansatz for the thresholds, we compute ⟨*s*⟩_*i*_ and obtain *w*_*i*_, which is used to derive the new set of thresholds.

In the case of the mixed population with ON and OFF cells, the optimal solution is one where the ON and OFF responses do not overlap; thus, there is no correlation between them. Therefore, we can treat each subpopulation separately, and in identical manner to the purely homogeneous case. The optimal weights and thresholds are identical to the homogeneous population population, with the exception of the constant term:

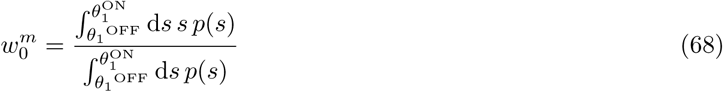

where *θ*_1_^OFF^ denotes the largest OFF threshold and *θ*_1_^ON^ denotes the smallest ON threshold in the population.

### Thresholds when optimizing the linear readout without noise

Now we consider the case of large *N* (for any mixture of ON and OFF cells) to derive the thresholds in the asymptotic limit where the threshold intervals (differences between neighboring thresholds) are small. We use a first order expansion of the stimulus distribution *p*(*s*) around each threshold *θ*_*j*_ in the expressions for ⟨*s*⟩_*j*_.

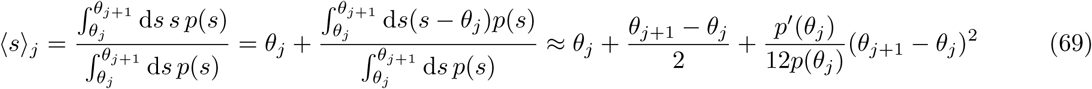

and similarly,

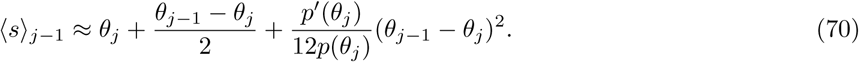

Combining Eq. 69 and Eq. 70 into Eq. 66, yields

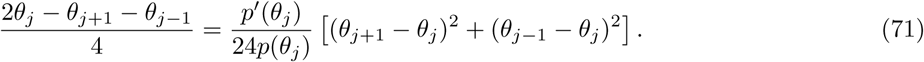

Taking the continuous limit so that *j* maps onto *x* with *j* = 1 corresponding to *x* = 0, *j* = *N* corresponding to *x* = 1, and d*x* = 1*/N*, we can write

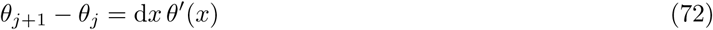

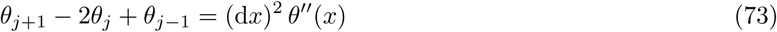

turning Eq. 71 into:

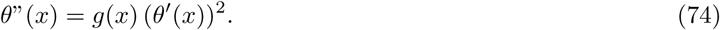

We can further define:

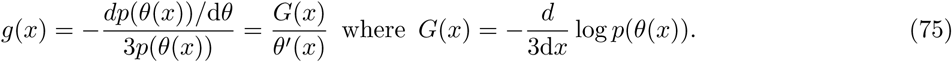

Denoting *y*(*x*) = *θ*′(*x*) gives the differential equation

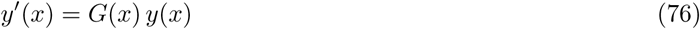

which has the solution

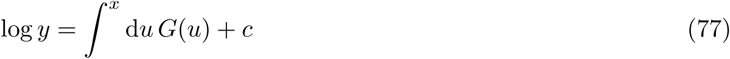

and consequently we obtain the differential equation

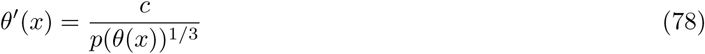

where *c* is a constant. This can be inverted into

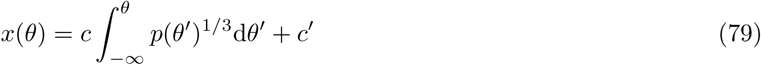

where *x* = *i/N* is the threshold index. We can determine the constants *c* and *c*′ from the boundary conditions:

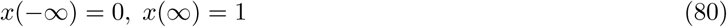

such that (as Eq. 8 in the Results section),

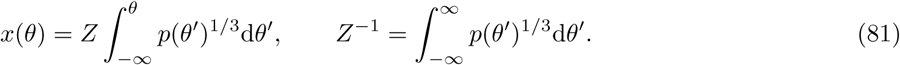

Inverting this relationship, we can obtain the threshold distribution *θ*(*x*) as a function of the index *x* = *i/N*. An expression for the optimal thresholds for the Laplace distribution: *p*(*s*) = 1*/*2*e*^−|*s*|^ is provided in Eq. 9 in the Results section.

### Optimal linear readout with noise: homogeneous population

For convenience, we normalize the linear readout

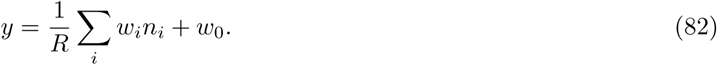

The error can be written as before (Eq. 60) with different correlations

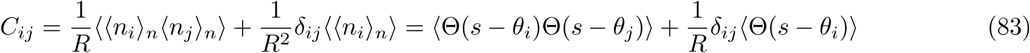

If we define, as before:

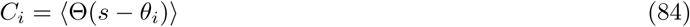

then for ⟨Θ(*s* − *θ*_*i*_)Θ(*s* − *θ*_*j*_) ⟩ = ⟨Θ(*s* − max(*θ*_*i*_, *θ*_*j*_)), and for *i* ≥ *j*:

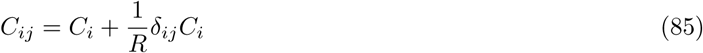

and

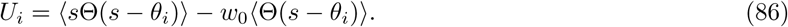

Optimizing with respect to the weights:

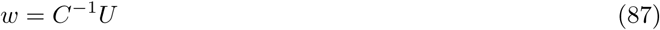

and

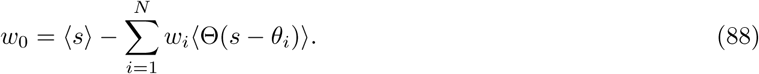

Optimizing with respect to the thresholds:

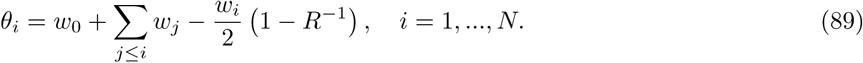

To solve these equations numerically, we implement an iterative procedure that rapidly converges to the optimal solution: starting from an ansatz for the thresholds, we compute *C* and *U* and obtain *w* from Eq. 87, which is used to derive the new set of thresholds.

### Thresholds when optimizing the linear readout with noise: homogeneous population

We provide an expression for the optimal thresholds for the general Laplace distribution:

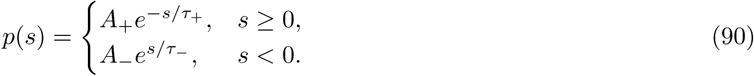

The symmetric Laplace distribution is one example, *p*(*s*) = 1*/*2*e*^−|*s*|^, with *A*_+_ = *A*_−_ = 1*/*2 and *τ*_+_ = *τ*_−_ = 1. In the limit of large population size *N*, we again derive the thresholds in the asymptotic limit where the threshold intervals are small. Assuming *θ*_1_ < 0,

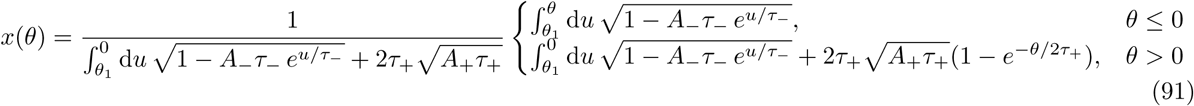

and assuming |*θ*_*1*_ | is large so that 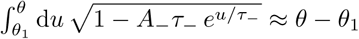, we can approximate

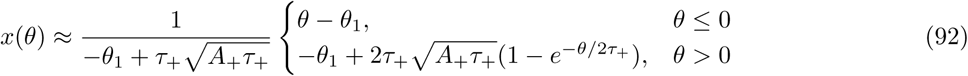

inverting this relationship, the optimal thresholds are:

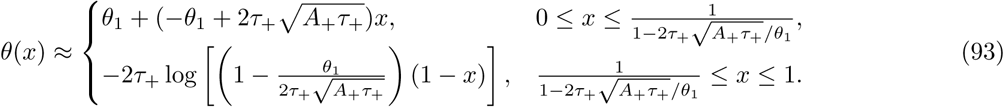

To fully determine the optimal thresholds, this requires knowledge of the first threshold, *θ*_1_. In the asymptotic limit, where the thresholds *θ*_*i*_ are close to each other, again expanding *p*(*s*) around each threshold, we derive

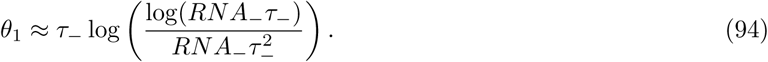

### Optimal linear readout with noise: mixed ON-OFF population

So far we have not explicitly treated the ON and OFF populations separately, because both when maximizing the mutual information for all noise levels, and minimizing the MSE in the limit of no noise, the performance and optimal thresholds were the same for all populations independent of the ON/OFF mixture. Now, we must treat the two populations separately.

Assume we have *m* ON cells and *N* − *m* OFF cells. We order the thresholds in the following manner (since non-overlapping ON and OFF cells is the optimal solution),

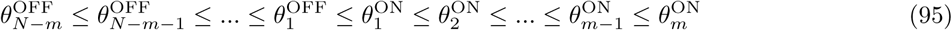

so that we can proceed in the same manner for each subpopulation as for the homogeneous population. The readout can be written as

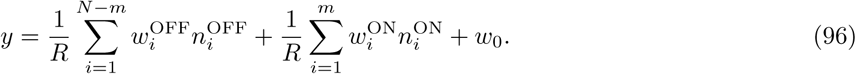

The error then is (assuming the optimal ON and OFF thresholds do not overlap – so that the ON-OFF cross-correlation is zero):

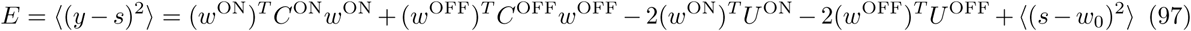

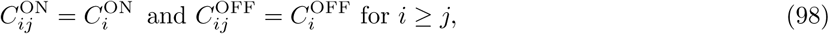

and

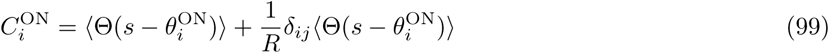

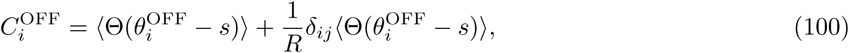

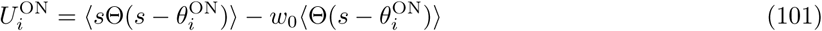

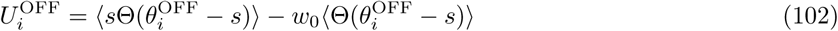

Optimizing with respect to the weights we get very similar expressions for each subpopulation (ON and OFF) as for the homogeneous population:

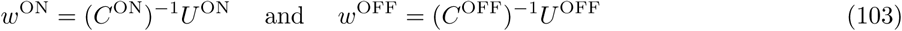

and optimizing the thresholds:

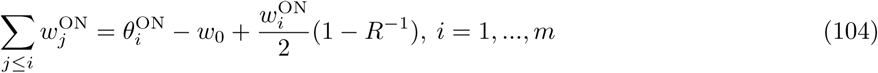

for the ON cells, and similarly for the OFF:

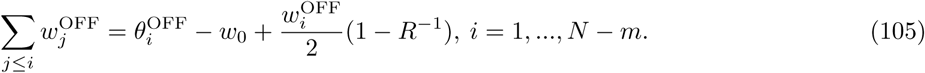

The difference from the homogeneous population is in the constant term:

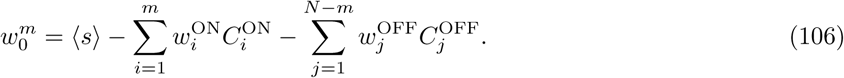

### Thresholds when optimizing the linear readout with noise: mixed ON-OFF population

We proceed in a similar fashion as with the homogeneous population to obtain the approximation in the case of large *N* : Let *f* ^ON^ = *m/N* be the fraction of ON cells and *f* ^OFF^ = (*N* − *m*)*/N* be the fraction of OFF cells in the population. We remap the thresholds, so that in the continuum limit 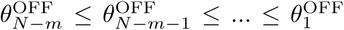 becomes *θ*^OFF^ (*x*^OFF^) and 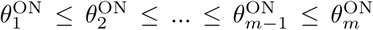 becomes *θ*^ON^ (*x*^ON^). Thus, the threshold index *x* = *i/N* ∈ [0, 1] for the homogeneous population becomes *x*^ON^ = *i/m* ∈ [0, *f* ^ON^] and *i* = 1, 2, …, *m* being the indices of the ON cels, and *x*^OFF^ = *i/*(*N* − *m*) ∈ [0, *f* ^OFF^] and *i* = 1, 2, …, *N* − *m* being the indices of the OFF cells. Figure 8 illustrates the mapping.

**Figure 8.**
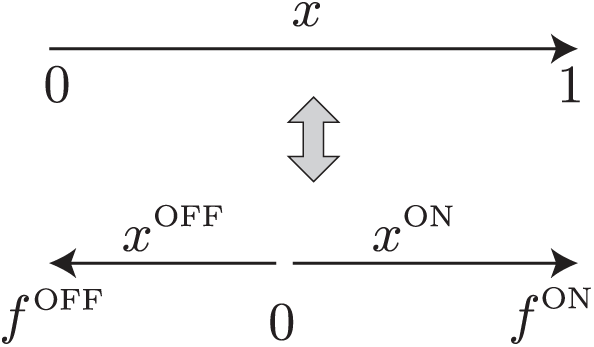
The mapping of the threshold indices from the homogeneous population with only ON cells to the mixed population with ON and OFF cells.

We provide an expression for the optimal thresholds for the general Laplace distribution (Eq. 90), and for a population that is unbalanced and has more ON cells, *f* ^ON^ *> f* ^OFF^. If 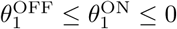, for the ON thresholds and weights the solution is similar to the case of the homogeneous population, i.e.

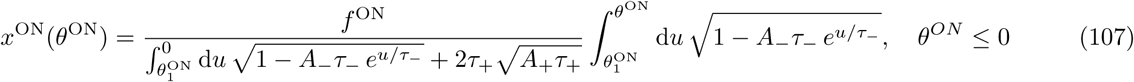

and

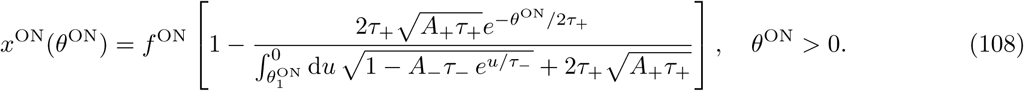

These expressions have to be inverted to obtain *θ*^ON^(*x*^ON^), which has to be done numerically. We proceed very similarly for the OFF cells. Namely, if 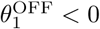 and assuming 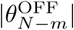 is large

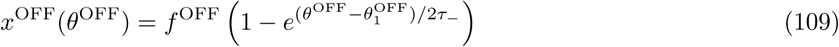

inverting this relationship is possible analytically

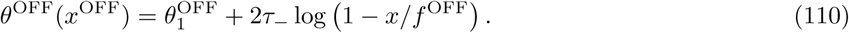

To fully determine the optimal thresholds, this requires knowledge of the first ON and OFF thresholds, 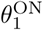 and 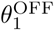.

When the population is mixed so that neither population dominates, the first ON and OFF thresholds are order 1. Assuming that they are close in stimulus space, so that 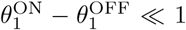, we can use the equations from optimizing the thresholds and weights to obtain the following equation which can be solved for 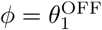:

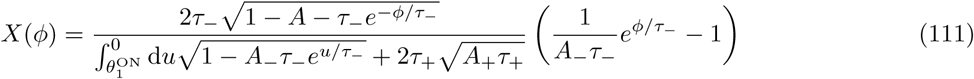

For the symmetric Laplace distribution with *A*_+_ = *A*_−_ = 1*/*2 and *τ*_+_ = *τ*_−_ = 1, the equation to solve for *ϕ* reduces to (Fig. 9):

**Figure 9.**
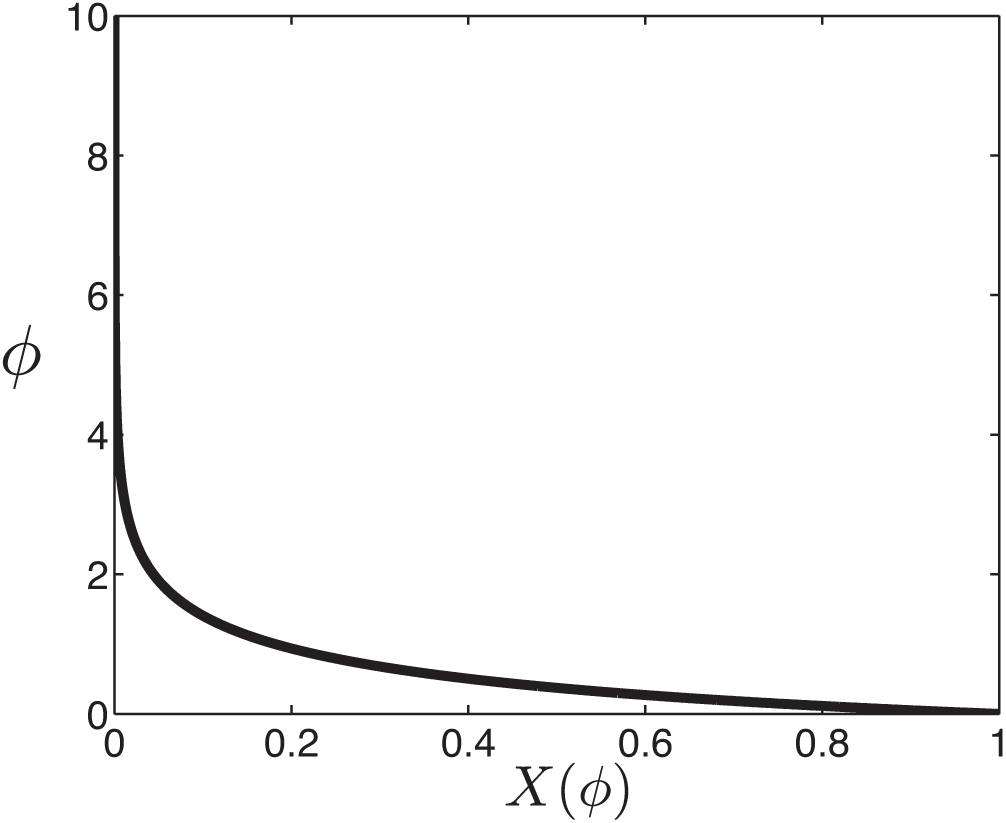
Determining the first thresholds for a mixed population of ON and OFF cells, 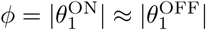 as a function of *X* = *f* ^OFF^*/f* ^ON^. For a symmetric Laplace distribution *p*(*s*) = 1*/*2*e*^−|*s*|^.

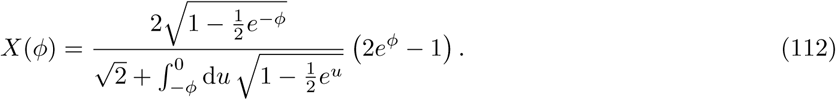

As shown in Figure 9, when there is an equal number of ON and OFF cells, *X* = 1 and 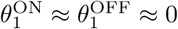. If there are 20% OFF cells and 80% ON cells in the population, then *X* = (1*/*5)*/*(4*/*5) = 1*/*4, and the first thresholds of each subpopulation are 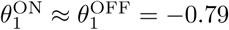. In the Results section we also considered asymmetric stimulus distributions where we varied the negative-to-positive bias *τ*_−_*/τ*_+_ and derived the solutions in a similar manner (Fig. 5).

### Deriving the stimulus distribution from measured ORN thresholds

From the study of Si and colleagues we extracted the distribution of measured thresholds (referred to as *EC*_50_ values) [37]. The cumulative distribution of the inverse of thresholds is

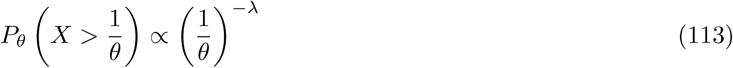

where *λ* = 0.42 (Fig. 6B). This enables us to derive the probability density function of the inverse of thresholds (Fig. 6C)

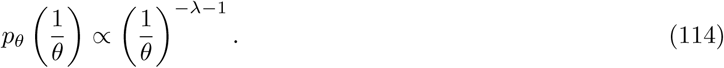

This distribution has a cut-off of *θ*_*c*_ = 4.22 · 10^4^ as reported in [37]. From this, we can derive the distribution of measured thresholds

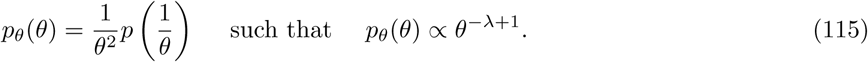

Next, we assume that these measured thresholds implement an optimal code first under the infomax criterion. Now, using the equation for the cumulative distribution of optimal thresholds in the large population limit, 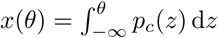, we can derive the stimulus distribution of odorant concentrations, *p*_*c*_,

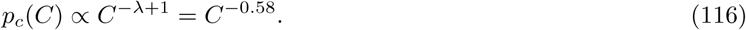

However, if we assume that these measured thresholds implement an optimal code under the criterion of minimizing the mean squared error of the optimal linear decoder, 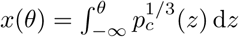, then the stimulus distribution of odorant concentrations, *p*_*c*_, is

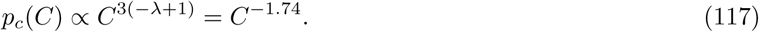

These are both shown in Fig. 6D.

## Supporting information

**S1 Figure. Binary neurons with spontaneous firing rate and Poisson noise**. A framework with binary neurons that have two firing rate levels, *r* if the stimulus is smaller (bigger) than a threshold, and *R* if the stimulus is bigger (smaller) than a threshold for ON (OFF) cells. We compare two systems, left: one ON (red) and one OFF (blue) cells, and right, two ON cells, where the information is maximized by optimizing the cells thresholds.

**S2 Figure. Sigmoidal neurons with sub-Poisson experimentally measured noise**. Two sigmoidal nonlinearities for an ON cell (red) and an OFF cell (blue), describing the firing rate as a function of stimulus with the maximum expected spike count *R*, the gain *β*, and the threshold *θ*. The shaded curve denotes the Laplace stimulus probability distribution.

**S1 Text. Mutual Information for a system with two cells**. The mutual information for different noise models.

**S1 Table. Conditional probability matrix**. Conditional probability matrix *p*(*k*_1_, *k*_2_|*s*) for a mixed ON-OFF system.

**S2 Table. Conditional probability matrix**. Conditional probability matrix *p*(*k*_1_, *k*_2_|*s*) for a homogeneous ON-ON system.

**S3 Table. Mutual information for a two-cell system**. Mutual information for a two-cell system with spontaneous firing rate and Poisson noise.

**S4 Table. Mutual information for a two-cell system**. Mutual information for a two-cell system with empirically measured sub-Poisson noise from salamander retinal ganglion cells.

## Acknowledgments

All authors were supported by the NIH, the Gatsby Charitable Foundation and the Swartz Foundation. JG was supported by the Max Planck Society and a Burroughs-Wellcome Career Award at the Scientific Interface. JG thanks Shuai Shao for careful reading of the analytical calculations.

## Notes

#### Summary of Updates

Mathematical details of the theory have been included in the Methods section. A new section on applying the efficient coding theory in reverse has been added, which applies to olfactory receptor neurons in Drosophila larva.

